# Discovery of genes required for body axis and limb formation by global identification of retinoic acid regulated epigenetic marks

**DOI:** 10.1101/778191

**Authors:** Marie Berenguer, Karolin F. Meyer, Jun Yin, Gregg Duester

## Abstract

Identification of target genes that mediate required functions downstream of transcription factors is hampered by the large number of genes whose expression changes when the factor is removed from a specific tissue and the numerous binding sites for the factor in the genome. Retinoic acid (RA) regulates transcription via RA receptors bound to RA response elements (RAREs) of which there are thousands in vertebrate genomes. Here, we combined ChIP-seq for epigenetic marks and RNA-seq on trunk tissue from wild-type and *Aldh1a2*-/-embryos lacking RA synthesis that exhibit body axis and forelimb defects. We identified a relatively small number of genes with altered expression when RA is missing that also have nearby RA-regulated deposition of H3K27ac (gene activation mark) or H3K27me3 (gene repression mark) associated with conserved RAREs, suggesting they have important downstream functions. RA-regulated epigenetic marks were identified near RA target genes already known to be required for body axis and limb formation, thus validating our approach, plus many other candidate RA target genes were found. *Nr2f1*, *Nr2f2*, *Meis1*, and *Meis2* gene family members were identified by our approach, and double knockouts of each family demonstrated previously unknown requirements for body axis and/or limb formation. These findings demonstrate that our method for identifying RA-regulated epigenetic marks can be used to discover genes important for development.

## Introduction

Retinoic acid (RA) is generated from retinol by the sequential activities of retinol dehydrogenase 10 (RDH10) (Sandell et al. 2007) and aldehyde dehydrogenase 1A2 (ALDH1A2) (Niederreither et al. 1999; Mic et al. 2002). Knockout studies of these enzymes revealed an essential role for RA in many early developmental programs including those controlling hindbrain anteroposterior patterning, neuromesodermal progenitor (NMP) differentiation, spinal cord neurogenesis, somitogenesis, forelimb bud initiation, and heart anteroposterior patterning (Rhinn and Dolle 2012; Cunningham and Duester 2015). RA functions as a ligand for nuclear RA receptors (RARs) that bind DNA sequences known as RA response elements (RAREs) as a heterodimer complex with retinoid X receptors (RXRs) (Mark et al. 2006). Binding of RA to RAR alters the ability of RAREs to recruit nuclear receptor coactivators (NCOAs) that activate transcription or nuclear receptor corepressors (NCORs) that repress transcription (Kumar et al. 2016). Thus, RA functions are mediated by transcriptional activation or repression of key genes via RAREs.

Identification of RA-regulated genes that are required for development has been difficult as loss or gain of RA activity alters the mRNA levels of thousands of genes in various cell lines or animals, perhaps most being indirect targets of RA or regulated post-transcriptionally. As RA target genes are dependent upon RAREs, identification of RAREs by RAR-binding studies, cell line transfection assays, and enhancer reporter transgenes in mouse or zebrafish has been used to identify RA target genes that may be required for development, but progress is slow as each gene is analyzed separately (Cunningham and Duester 2015). Genomic RAR chromatin immunoprecipitation (ChIP-seq) studies on mouse embryoid bodies and F9 embryonal carcinoma cells reported ∼14,000 potential RAREs in the mouse genome (Moutier et al. 2012; Chatagnon et al. 2015), but it is unclear how many of these RAREs are required to regulate genes in any specific tissue, and many may not function in any tissue at any stage of development. Only a few RAREs have been shown to result in gene expression and developmental defects when subjected to deletion analysis in mouse, i.e. a RARE enhancer that activates *Hoxa1* in the hindbrain (Dupé et al. 1997), a RARE enhancer that activates *Cdx1* in the spinal cord (Houle et al. 2003), and a RARE that functions as a silencer to repress caudal *Fgf8* in the developing trunk (Kumar et al. 2016). In one additional case, a RARE described within intron 2 of *Tbx5* that was suggested to be required for activation of *Tbx5* in the forelimb field based on a mouse enhancer reporter transgene (Nishimoto et al. 2015) was found to be unnecessary for *Tbx5* activation and forelimb budding when subjected to CRISPR deletion analysis, suggesting *Tbx5* is not an RA target gene (Cunningham et al. 2018). Many DNA control elements (including RAREs) that exhibit appropriate tissue-specific expression in enhancer reporter transgene assays have been shown to not be required as an enhancer in vivo when deleted; this may be due to enhancer redundancy or because the control element is really not an enhancer but appeared to be when inserted as a transgene at a random location in the genome near a heterologous promoter (Duester 2019). Thus, additional methods are needed (preferably genome-wide) to locate functional RAREs in a particular tissue which can be used to identify new candidate RA target genes that are required for development.

Epigenetic studies have found that histone H3 K27 acetylation (H3K27ac) associates with gene activation and histone H3 K27 trimethylation (H3K27me3) associates with gene repression (Rada-Iglesias et al. 2011; Laugesen and Helin 2014). We suggest that genes possessing nearby H3K27ac and H3K27me3 marks that are altered by loss of RA may point to direct transcriptional targets of RA (either activated or repressed) that are excellent candidates for performing functions downstream of RA. Here, we performed genomic ChIP-seq (H3K27ac and H3K27me3) and RNA-seq studies on E8.5 mouse embryonic trunks from wild-type and *Aldh1a2*-/-mouse embryos lacking RA synthesis to globally identify RA target genes for embryonic trunk. Candidate targets are defined as genes whose mRNA levels are decreased or increased by genetic loss of RA that also have nearby RA-regulated epigenetic marks associated with conserved RAREs, suggesting they have important downstream functions. This approach was able to identify many previously reported RA target genes known to control embryonic trunk development (including all three known RA target genes from RARE knockout studies: *Hoxa1*, *Cdx1*, and *Fgf8*), plus we identified numerous new candidate RA target genes that may control trunk development. CRISPR knockout studies on several of these new candidate RA target genes validated them as being required for body axis and/or limb formation. Our approach is generally applicable to determine tissue-specific target genes for any transcriptional regulator that has a knockout available.

## Results

### Comparison of RNA-seq and H3K27ac/H3K27me3 ChIP-seq for *Aldh1a2*-/-trunk tissue

Embryonic trunks were obtained from E8.5 embryos dissected to remove the head (including the pharyngeal region and anterior hindbrain), heart, and caudal tissue below the most recently formed somite as previously described (Kumar and Duester 2014). We performed RNA-seq analysis comparing E8.5 trunk tissue from wild-type embryos and *Aldh1a2*-/-embryos that lack the ability to produce RA (Mic et al. 2002). This analysis identified 4298 genes whose mRNA levels in trunk tissue are significantly decreased or increased when RA is absent (FPKM>0.5; a cut-off of log2 <-0.85 or >0.85 was employed to include *Sox2* known to be activated by RA; data available at GEO under accession number GSE131584).

We performed ChIP-seq analysis for H3K27ac and H3K27me3 epigenetic marks comparing E8.5 trunk tissue from wild-type and *Aldh1a2*-/-embryos isolated as described above for RNA-seq. This analysis identified 314 RA-regulated ChIP-seq peaks for H3K27ac located within or near 214 genes (i.e. the genes with the nearest annotated promoters) using a log2 cut-off of <-0.51 or >0.51 to include a RA-regulated peak near *Sox2* known to be activated by RA (Ribes et al. 2009; Cunningham et al. 2015). We identified 262 RA-regulated peaks for H3K27me3 located within or near 141 genes (i.e. the genes with nearest annotated promoters) using a log2 cut-off of <-0.47 or >0.47 to include a RA-regulated peak near *Fst* known to be repressed by RA (Cunningham et al. 2016); all ChIP-seq data available at GEO under accession number GSE131624. Thus, we found a much smaller number of RA-regulated ChIP-seq peaks for H3K27ac/H3K27me3 compared to the very large number of genes found to have altered mRNA levels with RNA-seq.

In order to identify genes that are good candidates for being transcriptionally activated or repressed by RA (RA target genes), we compared our ChIP-seq and RNA-seq results to identify RA-regulated ChIP-seq peaks where nearby genes have significant changes in expression in wild-type vs *Aldh1a2*-/-based on RNA-seq. We found 73 RA-regulated peaks for H3K27ac near 63 genes with significant changes in expression when RA is lost (Table S1), plus 46 RA-regulated peaks for H3K27me3 near 41 genes with significant changes in expression when RA is lost (Table S2). As some genes have more than one nearby RA-regulated peak for H3K27ac or H3K27me3, and some genes have nearby RA-regulated peaks for both H3K27ac and H3K27me3 (*Rarb*, *Dhrs3*, *Fgf8*, *Cdx2*, *Fst*, *Meis1*, *Meis2*, *Nr2f2*, *Foxp4*, *Ptprs*, and *Zfhx4*), a total of 93 RA-regulated genes have nearby RA-regulated peaks for H3K27ac and/or H3K27me3 when RA is lost, thus identifying them as candidate RA target genes for trunk development (Tables S1-S2; Fig. S1A).

**Table 1.**
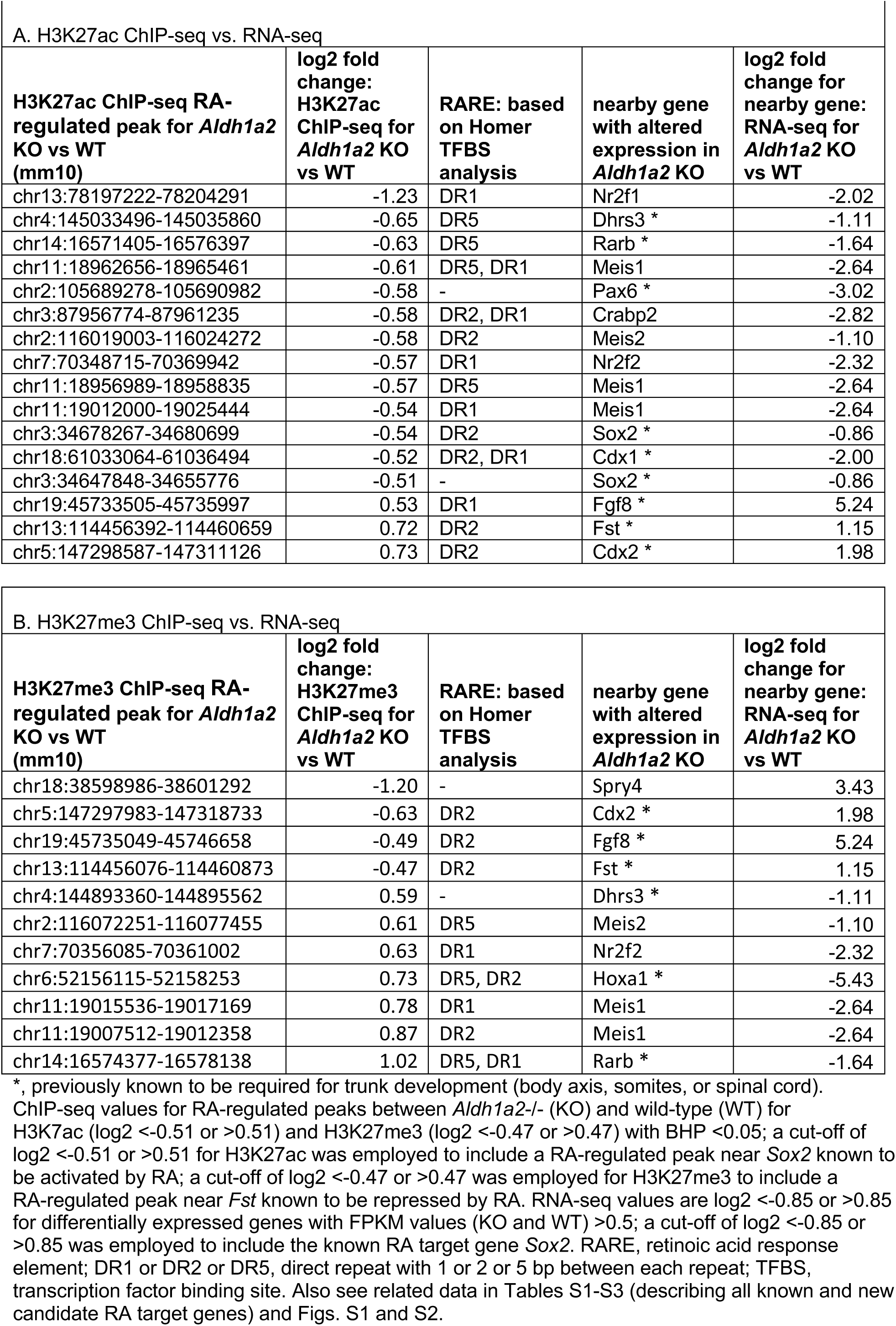
Comparison of ChIP-seq and RNA-seq for Aldh1a2-/-vs wild-type E8.5 trunk tissue showing identification of previously known RA target genes needed for trunk development plus some of the additional candidate RA target genes we identified here.

| A. H3K27ac ChIP-seq vs. RNA-seq |  |  |  |  |
| --- | --- | --- | --- | --- |
| H3K27ac ChIP-seq RA-regulated peak for *Aldh1a2* KO vs WT (mm10) | log2 fold change: H3K27ac ChIP-seq for *Aldh1a2* KO vs WT | RARE: based on Homer TFBS analysis | nearby gene with altered expression in *Aldh1a2* KO | log2 fold change for nearby gene: RNA-seq for *Aldh1a2* KO vs WT |
| chr13:78197222-78204291 | -1.23 | DR1 | Nr2f1 | -2.02 |
| chr4:145033496-145035860 | -0.65 | DR5 | Dhrs3 * | -1.11 |
| chr14:16571405-16576397 | -0.63 | DR5 | Rarb * | -1.64 |
| chr11:18962656-18965461 | -0.61 | DR5, DR1 | Meis1 | -2.64 |
| chr2:105689278-105690982 | -0.58 | - | Pax6 * | -3.02 |
| chr3:87956774-87961235 | -0.58 | DR2, DR1 | Crabp2 | -2.82 |
| chr2:116019003-116024272 | -0.58 | DR2 | Meis2 | -1.10 |
| chr7:70348715-70369942 | -0.57 | DR1 | Nr2f2 | -2.32 |
| chr11:18956989-18958835 | -0.57 | DR5 | Meis1 | -2.64 |
| chr11:19012000-19025444 | -0.54 | DR1 | Meis1 | -2.64 |
| chr3:34678267-34680699 | -0.54 | DR2 | Sox2 * | -0.86 |
| chr18:61033064-61036494 | -0.52 | DR2, DR1 | Cdx1 * | -2.00 |
| chr3:34647848-34655776 | -0.51 | - | Sox2 * | -0.86 |
| chr19:45733505-45735997 | 0.53 | DR1 | Fgf8 * | 5.24 |
| chr13:114456392-114460659 | 0.72 | DR2 | Fst * | 1.15 |
| chr5:147298587-147311126 | 0.73 | DR2 | Cdx2 * | 1.98 |

| B. H3K27me3 ChIP-seq vs. RNA-seq |  |  |  |  |
| --- | --- | --- | --- | --- |
| H3K27me3 ChIP-seq RA-regulated peak for *Aldh1a2* KO vs WT (mm10) | log2 fold change: H3K27me3 ChIP-seq for *Aldh1a2* KO vs WT | RARE: based on Homer TFBS analysis | nearby gene with altered expression in *Aldh1a2* KO | log2 fold change for nearby gene: RNA-seq for *Aldh1a2* KO vs WT |
| chr18:38598986-38601292 | -1.20 | - | Spry4 | 3.43 |
| chr5:147297983-147318733 | -0.63 | DR2 | Cdx2 * | 1.98 |
| chr19:45735049-45746658 | -0.49 | DR2 | Fgf8 * | 5.24 |
| chr13:114456076-114460873 | -0.47 | DR2 | Fst * | 1.15 |
| chr4:144893360-144895562 | 0.59 | - | Dhrs3 * | -1.11 |
| chr2:116072251-116077455 | 0.61 | DR5 | Meis2 | -1.10 |
| chr7:70356085-70361002 | 0.63 | DR1 | Nr2f2 | -2.32 |
| chr6:52156115-52158253 | 0.73 | DR5, DR2 | Hoxa1 * | -5.43 |
| chr11:19015536-19017169 | 0.78 | DR1 | Meis1 | -2.64 |
| chr11:19007512-19012358 | 0.87 | DR2 | Meis1 | -2.64 |
| chr14:16574377-16578138 | 1.02 | DR5, DR1 | Rarb * | -1.64 |

\*, previously known to be required for trunk development (body axis, somites, or spinal cord).

**Table 2.**
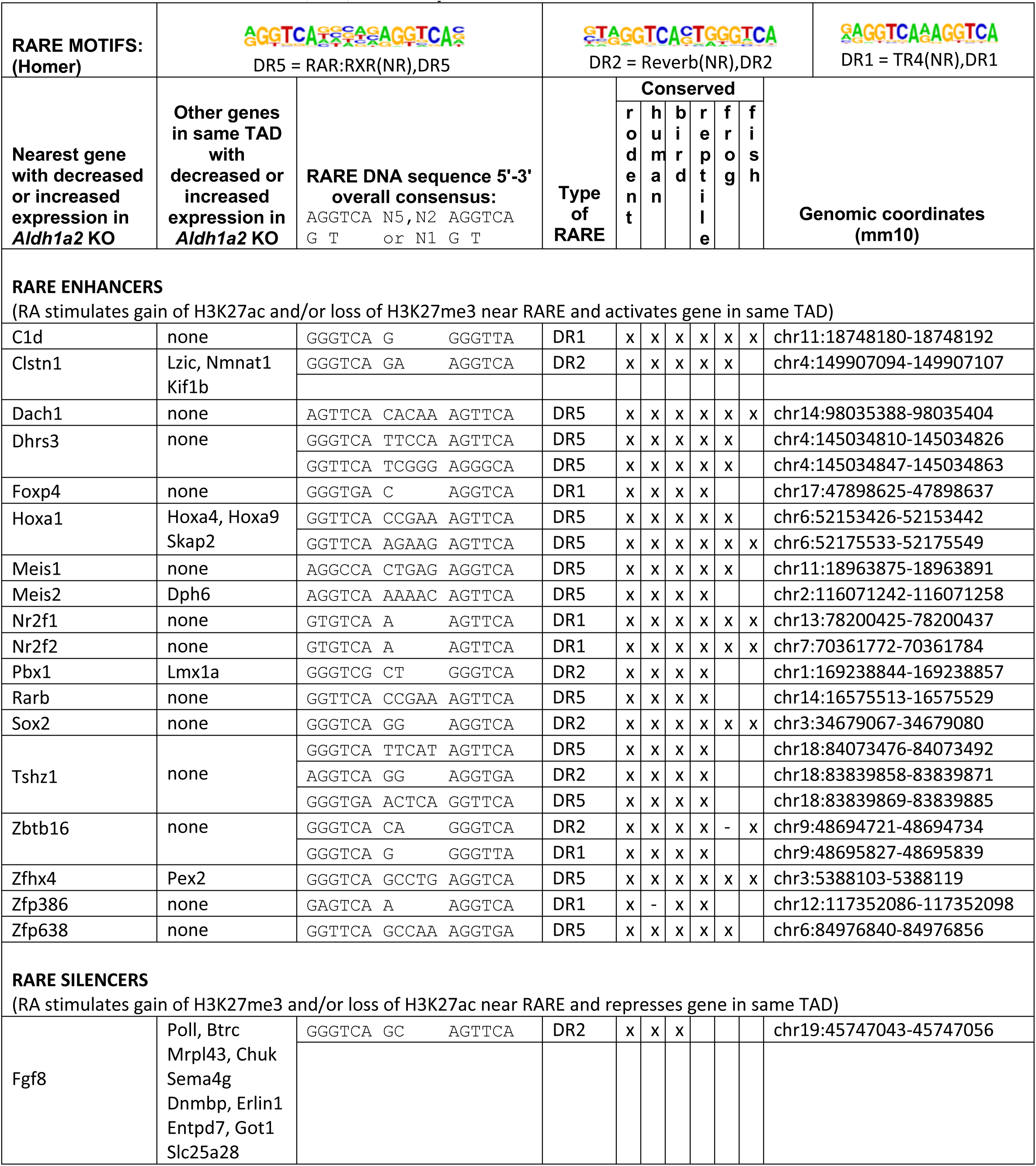
DNA sequences of highly conserved RAREs located in RA-regulated ChIP-seq peaks for H3K27ac or H3K27me3 near all RA-regulated genes in same TAD. RAREs shown here are conserved from mouse to bird, reptile, frog, or fish. RAREs contain no more than one mismatch to Homer consensus DR5, DR2, or DR1 RARE motifs shown here; DR, direct repeat.

Among the 93 candidate RA target genes for trunk development identified with our approach are included many examples of genes previously reported to be regulated by RA in the trunk based on studies of *Aldh1a2*-/-embryos (Niederreither and Dolle 2008; Cunningham and Duester 2015) or RA-treated NMPs (Cunningham et al. 2016); this includes *Hoxa1*, *Cdx1*, *Rarb*, *Crabp2*, *Sox2*, *Dhrs3*, and *Pax6* whose expression is increased by RA, plus *Fgf8*, *Cdx2*, and *Fst* whose expression is decreased by RA (Table 1). H3K27ac peaks near *Cdx1, Rarb*, *Crabp2*, *Sox2*, *Dhrs3*, and *Pax6* are reduced in *Aldh1a2*-/-trunk consistent with these being RA-activated genes, whereas H3K27ac peaks near *Fgf8*, *Cdx2*, and *Fst* are increased in *Aldh1a2*-/-consistent with these being genes repressed by RA. Conversely, H3K27me3 peaks near *Fgf8*, *Cdx2*, and *Fst* are decreased in *Aldh1a2*-/-, whereas H3K27me3 peaks near *Rarb*, *Hoxa1*, and *Dhrs3* are increased in *Aldh1a2*-/-, consistent with the former being genes repressed by RA and the latter being genes activated by RA (Table 1). In addition to these 10 well-established RA target genes that are required for trunk development, we identified 83 additional genes that our findings indicate are candidate RA target genes for trunk, including *Nr2f1, Nr2f2, Meis1, Meis2,* and *Spry4* that were further examined here (Table 1); differential expression of these genes in E8.5 wild-type vs *Aldh1a2*-/-trunk was validated by qRT-PCR (Fig. S2). As our approach identified many known trunk RA target genes, it is a reliable approach for identifying new candidate RA target genes required for trunk development.

### Identification of RAREs associated with RA-regulated deposition of H3K27ac or H3K27me epigenetic marks

As RA target genes need to be associated with a RARE, the DNA sequences within the RA-regulated H3K27ac/H3K27me3 ChIP-seq peaks we found near our list of 93 RA-regulated genes were searched for RARE sequences using the Homer transcription factor binding site program for the mm10 genome; we searched for three types of RAREs including those with a 6 bp direct repeat separated by either 5 bp (DR5), 2 bp (DR2), or 1 bp (DR1) (Cunningham and Duester 2015), and the presence or absence of RAREs is summarized (Tables S1 and S2). We found that 46 of these 93 genes contained at least one RARE in their nearby RA-regulated H3K27ac and/or H3K27me3 ChIP-seq peaks, thus narrowing down our list of candidate RA target genes to 49% of the genes originally identified. Our approach identified the three RAREs previously shown to have required functions during trunk development in vivo by knockout studies (RAREs for *Hoxa1*, *Cdx1*, *Fgf8*) plus several RAREs associated with known RA-regulated genes in the E8.5 trunk from *Aldh1a2*-/-studies (*Rarb*, *Crabp2*, *Sox2*, *Dhrs3*, *Cdx2*, *Fst*), thus validating our approach for identifying RA-regulated genes required for trunk development. The sequences of all the RAREs near these 46 RA target genes here are summarized; included are 65 RAREs near 34 RA-activated genes (we refer to these as RARE enhancers associated with increased H3K27ac and/or decreased H3K27me3 in the presence of RA) and 20 RAREs near 12 RA-repressed genes (we refer to these as RARE silencers associated with increased H3K27me3 and/or decreased H3K27ac in the presence of RA) (Table S3).

The results here provide evidence that many of the RA-regulated H3K27ac and H3K27me3 marks are associated with regulation of the nearest genes. However, it is possible that some H3K27ac and H3K27me3 RA-regulated peaks may be related to RA-regulated genes located further away in the same topologically associated domain (TAD). In order to address this issue, we assigned each RA-regulated H3K27ac and H3K27me3 peak to a TAD using the 3D Genome Browser (http://promoter.bx.psu.edu/hi-c/view.php); TAD analysis has not been performed on mouse E8.5 trunk tissue, but as TAD domains are similar between different tissues (Dixon et al. 2012) we used the TAD database for mouse ES cells which is the closest biologically relevant database in the 3D Genome Browser. Then the genes in each TAD containing an RA-regulated peak were searched in our RNA-seq database to identify genes whose mRNA levels are decreased or increased when RA is lost, and if at least one gene was found we determined whether a RARE is present in the ChIP-seq peak. This analysis resulted in the identification of 82 additional RARE enhancers near RA-activated genes, and 40 additional RARE silencers near RA-repressed genes, where the gene is not the gene nearest to the RARE in the TAD; in some cases more than one RA-regulated gene was identified in a TAD (Table S3).

Up to now, *Fgf8* represents the only example of a gene that is directly repressed by RA at the transcriptional level as shown by developmental defects upon knockout of the RARE at −4.1 kb, and by the ability of this RARE to stimulate binding of NCOR and PRC2 plus deposition of H3K27me3 in an RA-dependent manner (Kumar and Duester 2014; Kumar et al. 2016). Here, in addition to *Fgf8*, we found many more candidates for genes repressed by RA in the trunk based on identification of nearby RARE silencers (Tables S3).

### Analysis of known RA target genes for trunk validates our approach for finding new targets

The RA-regulated H3K27ac and/or H3K27me3 peaks we identified near *Rarb*, *Crabp2*, *Hoxa1*, and *Cdx1* all overlap previously reported RAREs for these genes (Fig. 1). In the case of *Rarb*, the DR5 RARE in the 5’-untranslated region (Mendelsohn et al. 1991) overlaps RA-regulated peaks for both H3K27ac and H3K27me3, suggesting that this RARE in the presence of RA stimulates deposition of H3K27ac and removal of H3K27me3 during activation of *Rarb*; we identified a DR1 RARE in the 5’-noncoding region of *Rarb* within another RA-regulated H3K27me3 ChIP-seq peak (Fig. 1A). For *Crabp2*, two closely-spaced RAREs previously reported in the 5’-noncoding region (Durand et al. 1992) associate with RA-regulated peaks for H3K27ac, plus another RARE we identified in the 3’-noncoding region associates with changes in H3K27ac (Fig. 1B). For *Hoxa1*, the RARE located in the 3’-noncoding region is associated with RA-regulated peaks for both H3K27ac and H3K27me3, plus another RARE we identified in the 3’-untranslated region is associated with RA-regulated peaks for H3K27me3 (Fig. 1C); importantly, knockout studies on the *Hoxa1* RARE in the 3’-noncoding region demonstrated that it is required *in vivo* for *Hoxa1* expression and normal development (Dupé et al. 1997). For *Cdx1*, two RAREs have been reported, one in the 5’-noncoding region that was shown by knockout studies to be required for *Cdx1* expression and body axis development (Houle et al. 2003), plus another RARE in intron 1 (Gaunt and Paul 2011). Both of these *Cdx1* RAREs are overlapped by RA-regulated peaks for both H3K27ac and H3K27me3 (Fig. 1D). These findings demonstrate that our approach can identify genes that are already known to be transcriptionally activated by RA via a RARE and required for development.

**Fig. 1.**
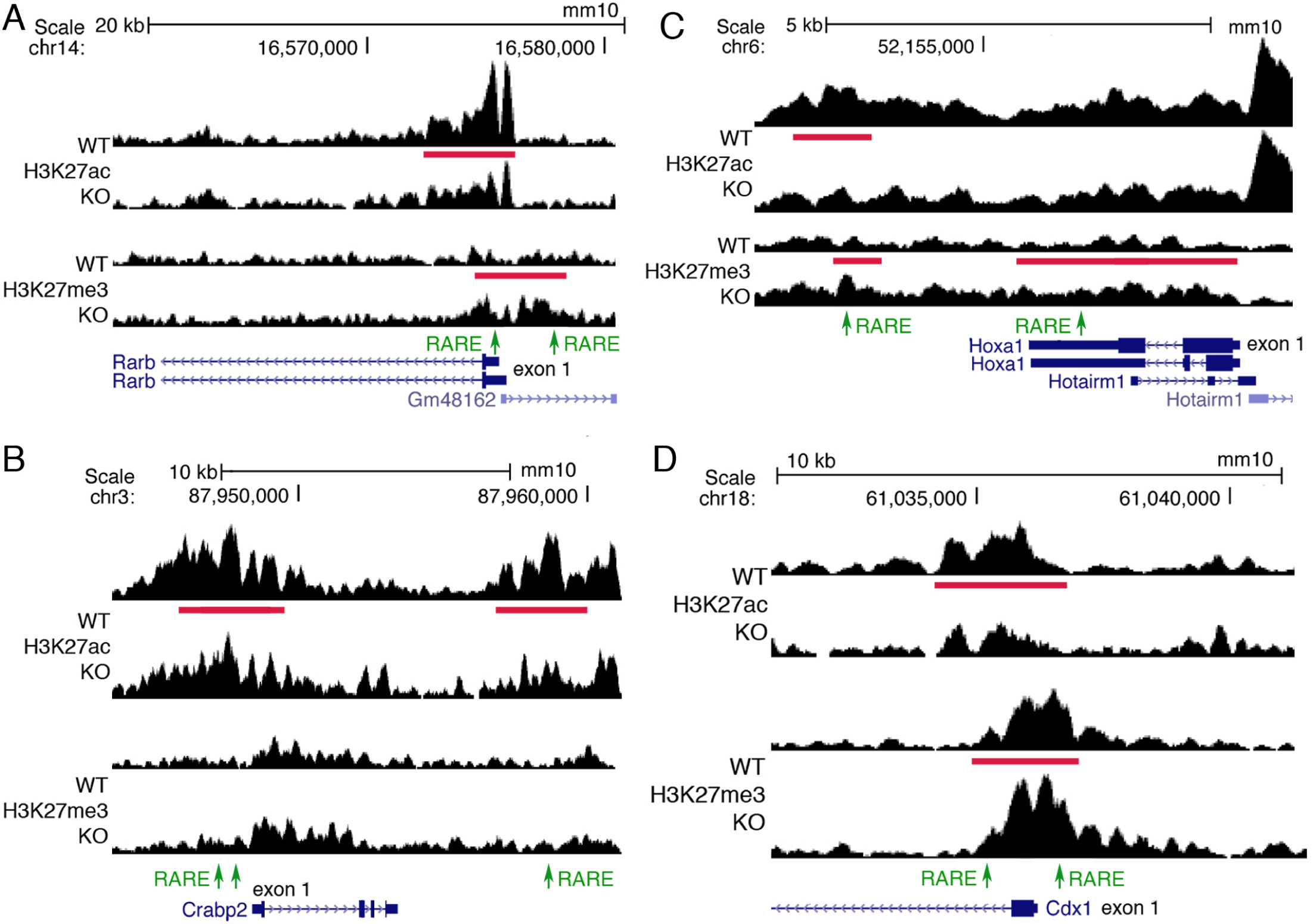
ChIP-seq findings for *Rarb*, *Crabp2*, *Hoxa1*, and *Cdx1* showing that RA-regulated peaks for H3K27ac and H3K7me3 are located near known RARE enhancers. (A) Shown for *Rarb* are RA-regulated ChIP-seq peaks for H3K27ac and H3K27me3 (red bars) when RA is lost in E8.5 trunk comparing wild-type (WT) vs *Aldh1a2*-/-(KO) as well as RAREs (green). A RARE in the 5’-untranslated region is known to function as an RA-dependent enhancer in mouse transgene studies (ref. 21); here, H3K27ac is decreased and H3K27me3 increased near the native RARE when RA is lost in trunk tissue, supporting its function as a RARE enhancer in vivo. We also found a RARE in the 5’-noncoding region of *Rarb* within an H3K27me3 ChIP-seq peak that is increased when RA is lost. (B) RA-regulated peaks for H3K27ac and RAREs are shown for *Crabp2*. The two RAREs in the 5’-noncoding region were previously shown to function as RA-dependent enhancers in cell line studies (ref. 22). Our epigenetic studies also identified another RARE enhancer in the 3’-noncoding region. (C) RA-regulated peaks for H3K27ac and/or H3K27me3 and RAREs are shown for *Hoxa1*. Knockout studies in mouse embryos have shown that the RARE in the 3’-noncoding region is essential for hindbrain *Hoxa1* expression and development (ref. 10). (D) RA-regulated peaks for H3K27ac and H3K27me3 and RAREs are shown for *Cdx1*. Knockout studies in mouse embryos have shown that the RARE in the 5’-noncoding region is essential for *Cdx1* expression and body axis development (ref. 11). RA-regulated peaks in the genome browser view shown here and elsewhere are for one replicate, with the other replicate showing a similar result.

### Identification of RA-regulated epigenetic marks and RAREs near RA-regulated genes known to control neuromesodermal progenitors

Ingenuity Pathway Analysis (IPA) of our list of 93 RA target genes shows enrichment for the pathway “development of body trunk”, including *Sox2*, *Cdx2*, and *Fgf8* known to be required for neuromesodermal progenitor (NMP) function during trunk development (Fig. S1B). NMPs are bipotential progenitor cells in the caudal region co-expressing *Sox2* and *T/Bra* that undergo balanced differentiation to either spinal cord neuroectoderm or presomitic mesoderm to generate the post-cranial body axis (Wilson et al. 2009; Kondoh and Takemoto 2012; Henrique et al. 2015; Amin et al. 2016; Kimelman 2016; Gouti et al. 2017; Koch et al. 2017; Edri et al. 2019). NMPs are first observed in mouse embryos at about E8.0 near the node and caudal lateral epiblast lying on each side of the primitive streak (Olivera-Martinez et al. 2012; Tsakiridis et al. 2014; Garriock et al. 2015). Caudal Wnt and FGF signals are required to establish and maintain NMPs (Naiche et al. 2011; Takemoto et al. 2011; Martin and Kimelman 2012; Olivera-Martinez et al. 2012; Jurberg et al. 2014; Garriock et al. 2015; Wymeersch et al. 2016). *Cdx2* is required for establishment of NMPs (Amin et al. 2016). During development, RA is first produced at E7.5 in presomitic mesoderm expressing *Aldh1a2* to generate an anteroposterior gradient of RA with high activity in the trunk and low activity caudally (Cunningham and Duester 2015). Loss of RA does not prevent establishment or maintenance of NMPs, but does result in unbalanced differentiation of NMPs, with decreased caudal *Sox2* expression and decreased appearance of neural progenitors, plus increased caudal *Fgf8* expression and increased appearance of mesodermal progenitors and small somites due to encroachment of caudal *Fgf8* expression into the trunk where it reduces epithelial condensation of presomitic mesoderm needed to form somites (Diez del Corral et al. 2003; Patel et al. 2013; Cunningham et al. 2015; Cunningham et al. 2016). *Cdx2* expression is increased when RA is lost in *Aldh1a2*-/-embryos (Zhao and Duester 2009).

Here, when RA is lost we observed RA-regulated H3K27ac and/or H3K27me3 peaks near several genes required for NMP function that show decreased (*Sox2*) or increased (*Fgf8* and *Cdx2*) expression (Fig. 2A-C). Most of these RA-regulated peaks contain RAREs, providing evidence that *Sox2*, *Fgf8*, and *Cdx2* are direct RA target genes (Table S3). For *Sox2*, we observed two RA-regulated H3K27ac ChIP-seq peaks, but only the one in the 3’-noncoding region was found to have a RARE (Fig. 2A). In the case of *Fgf8*, previous studies reporting knockout of the RARE located in the 5’-noncoding region at −4.1 kb resulted in increased caudal *Fgf8* expression and a small somite phenotype (although the defect is not as severe as for *Aldh1a2*-/-embryos), demonstrating that this RARE functions in vivo as a silencer by RA-dependent recruitment of nuclear receptor corepressors (Kumar et al. 2016); RARE redundancy may explain the milder phenotype as our approach suggests that *Fgf8* has two additional candidate RARE silencers (Fig. 2B). RARE redundancy may be common as we observe that *Cdx2* has three candidate RARE silencers (Fig. 2C), and our overall analysis shows that many genes have more than one nearby RARE (Table S3). These findings provide evidence that RA controls NMP differentiation directly at the transcriptional level by activating *Sox2* and repressing *Fgf8* and *Cdx2* as progenitor cells progress from a caudal to a trunk location.

**Fig. 2.**
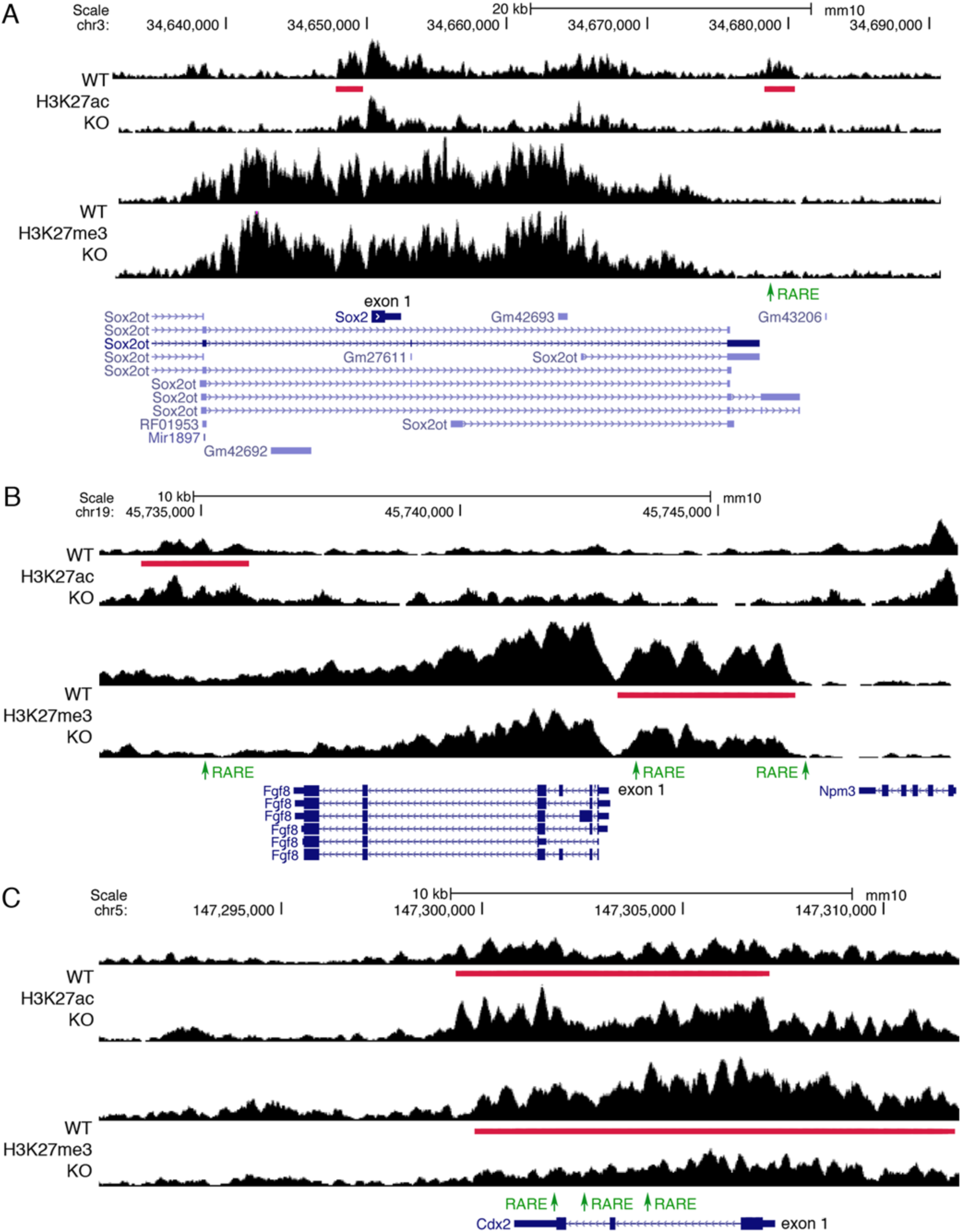
ChIP-seq findings identify RAREs near genes required for NMP function. (A) Two RA-regulated ChIP-seq peaks for H3K27ac (red bars) near *Sox2* are shown for trunk tissue from E8.5 wild-type (WT) vs *Aldh1a2*-/-(KO). A RARE (green) was found in the 3’-noncoding peak (but not the 5’-noncoding peak) suggesting it may function as a RARE enhancer as the H3K27ac peak is decreased when RA is lost. (B) Shown are RA-regulated ChIP-seq peaks for H3K27me3 and H3K27ac near *Fgf8*. In the 5’-noncoding region of *Fgf8* we found two RAREs on either end of the peak for H3K27me3 (repressive mark) that is decreased in KO, indicating they are candidate RARE silencers; the RARE furthest upstream in the 5’-noncoding region at −4.1 kb was shown by knockout studies to function as an RA-dependent RARE silencer required for caudal *Fgf8* repression and somitogenesis (ref. 7). We also found another RARE in the 3’-noncoding region of *Fgf8* that is another candidate for a RARE silencer as it is contained within a RA-regulated peak for H3K27ac (activating mark) that is increased when RA is lost. (C) *Cdx2* has a peak for H3K27ac that is increased and an overlapping peak for H3K27me3 that is decreased, along with three RAREs included within both peaks, indicating that all these RAREs are candidates for RARE silencers.

### Evidence for genes regulated indirectly by RA at the transcriptional level

Our studies show that many genes that are downregulated or upregulated following loss of RA are associated with RA-regulated peaks for H3K27ac or H3K27me3 (either nearby or in the same TAD) that do not contain RAREs (Tables S1-S2). Such genes may be indirectly activated or repressed by RA at the transcriptional level. In the case of *Pax6*, our results indicate that RA stimulates H3K27ac deposition in *Pax6* introns 2 and 6 that do not contain RAREs, with no other RA-regulated peaks in the same TAD (Fig. 3A). Previous studies identified an enhancer in *Pax6* intron 6 containing a SOXB1 binding site that is important for activation in the spinal cord (Oosterveen et al. 2013). Activation of *Pax6* in the spinal cord requires CDX proteins in the posterior-most neural tube, and CDX binding sites have been identified in *Pax6* intron 2 (Joshi et al. 2019); in addition to expression in the caudal progenitor zone, mouse *Cdx1* is expressed in the posterior neural plate where *Pax6* is activated, and this expression domain requires RA (Zhao and Duester 2009). Activation of *Pax6* requires that caudal FGF signaling be downregulated (Patel et al. 2013). Thus, although it is possible that our H3K27ac/H3K27me3 studies failed to identify an unknown RARE near *Pax6*, our findings suggest that the RA requirement for *Pax6* activation may operate through several indirect mechanisms due to the ability of RA to activate *Sox2* and *Cdx1*, and repress *Fgf8* (Figs. 1, 2).

**Fig. 3.**
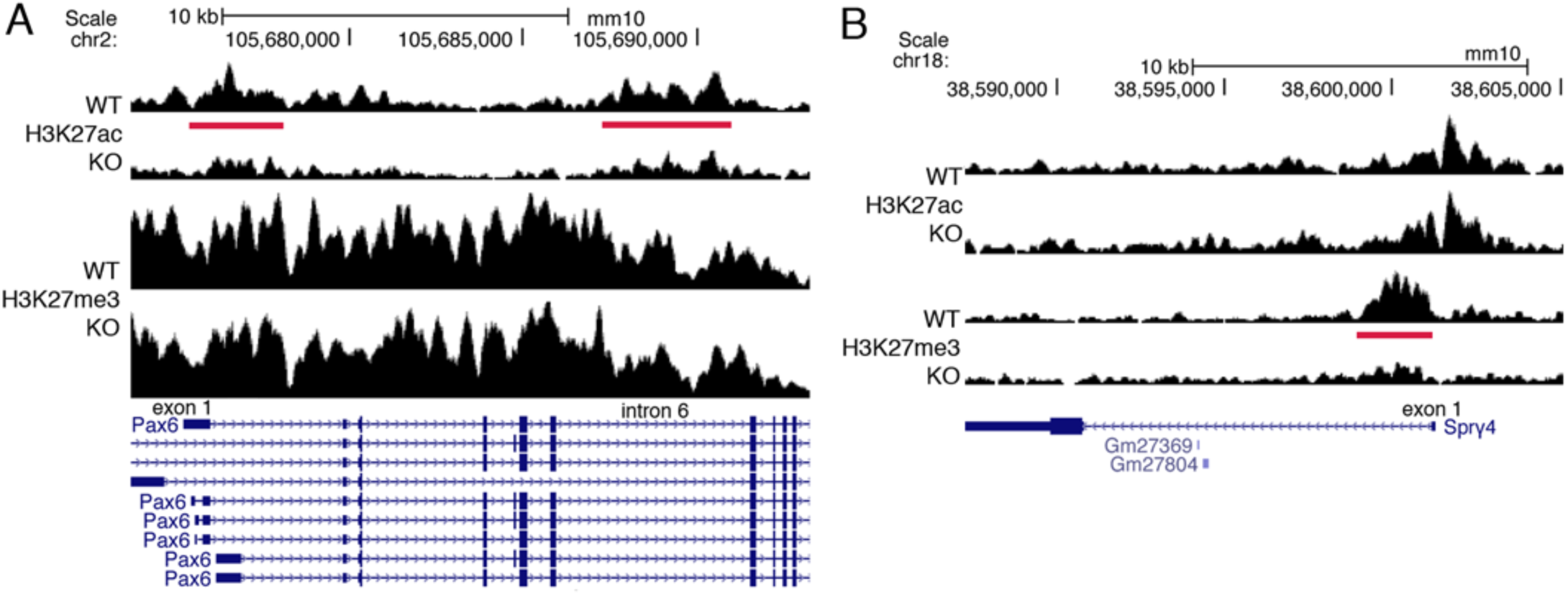
ChIP-seq findings for *Pax6* and *Spry4* that lack RARE enhancers or silencers. These genes are good candidates for being indirect transcriptional targets of RA as their RA-regulated ChIP-seq peaks do not contain RAREs. (A) *Pax6* has two RA-regulated peaks (red bars) for H3K27ac (decreased) when RA is lost in E8.5 trunk tissue from *Aldh1a2*-/-(KO) compared to wild-type (WT); these RA-regulated peaks do not contain RAREs suggesting that transcription of *Pax6* is indirectly activated by RA. (B) *Spry4* has an RA-regulated peak for H3K27me3 (decreased) when RA is lost with no associated RARE suggesting that transcription of *Spry4* is indirectly repressed by RA.

We observed that *Spry4* (shown here to be down-regulated by RA) does not have a RARE associated with its RA-regulated ChIP-seq peak for H3K27me3; no other RA-regulated peaks were found in its TAD (Fig. 3B). Many of the RA-regulated ChIP-seq peaks observed with our approach that do not contain RAREs may be indirect RA-regulated peaks that contain DNA binding sites for transcription factors other than RARs whose expression or activity is altered by loss of RA, thus resulting in changes for H3K27ac/H3K27me3 marks that are caused by the other transcription factors.

### Conservation of RAREs identified with our approach identifies candidate RA target genes

While it possible that some RAREs that are conserved only in mammals perform mammal-specific functions, the RAREs we found in mouse that are conserved to birds or lower provide further evidence that they are needed to regulate target genes. The candidate RARE enhancers and RARE silencers we identified here that are associated with RA-regulated epigenetic marks were searched for evolutionary conservation using the UCSC genome browser. Among the RAREs in which the nearest gene is RA-regulated we found 6 RAREs that are conserved from mouse to zebrafish, 11 conserved to frog (*X. tropicalis*), 18 conserved to reptile (lizard; painted turtle), 20 conserved to bird (chicken; turkey), 39 conserved to human, 65 conserved to rodent (rat), and 20 that are not conserved with rat (Table S3). The large number of RAREs (i.e. 20) conserved beyond mammals to bird, lizard, frog, or fish demonstrate that our approach is able to identify highly conserved RAREs that point to excellent candidate genes required for development. Among the additional RAREs we found located further away in the TAD from an RA-regulated gene we identified only 4 more RAREs conserved beyond mammals to bird, lizard, frog, or fish, thus bringing the total to 24 highly conserved RAREs (Table S3). Thus, most of the highly conserved RAREs we identified are located very close to an RA-regulated gene rather than further distant in the TAD. In addition, all these highly conserved RAREs are either identical to the RARE consensus or have only one mismatch. Here we summarize the 24 most highly conserved RAREs that point to 38 RA-regulated genes that may be required for development (Table 2).

As RAREs need to be bound by RAR in order to function, we examined previously reported RAR ChIP-seq databases for mouse embryoid bodies (Moutier et al. 2012) and mouse F9 embryonal carcinoma cells (Chatagnon et al. 2015) to determine if the highly conserved RAREs we identified are included in RAR-binding regions. We found that 19 of our 24 highly conserved RAREs are included in the RAR ChIP-seq peaks from at least one of those studies (Table S4).

Our list of best candidate RA target genes (Table 2) includes several for which knockout studies have already demonstrated required functions during trunk development, i.e. in RA signaling (*Rarb, Dhrs3*), body axis formation (*Hoxa1, Hoxa4, Hoxa9, Sox2, Fgf8, Pbx1, Tshz1, Zbtb16*), and foregut formation (*Foxp4*); mouse knockout data summarized by Mouse Genome Informatics (http://www.informatics.jax.org). This list also includes many genes for which knockout studies have either not been performed or knockouts resulted in no reported early developmental defects. This list of genes thus contains excellent new candidates that can be tested for function during trunk development by generating knockouts or double knockouts in the case of gene families.

### *Nr2f* and *Meis* gene families have nearby RA-regulated epigenetic marks associated with highly conserved RARE enhancers

We identified two gene families (*Nr2f* and *Meis*) where two family members have decreased expression when RA is lost and nearby RA-regulated peaks for H3K27ac or H3K27me3 containing RAREs. *Nr2f1* and *Nr2f2* encode orphan nuclear receptors NR2F1 and NR2F2 (previously known as COUP-TFI and COUP-TFII, respectively) that regulate transcription, although they have not been found to have endogenous ligands that control their activity (Lin et al. 2011). *Meis1* and *Meis2* encode transcription factors belonging to the TALE (three amino acid loop extension) family of homeodomain-containing proteins (Penkov et al. 2013; Schulte and Geerts 2019).

Previous studies suggested that *Nr2f* genes are activated by RA in *Ciona*, zebrafish, and mouse cell lines (Ishibashi et al. 2005; Zhuang and Gudas 2008; Laursen et al. 2013; Dohn et al. 2019). Here, *Nr2f1* and *Nr2f2* were both found to have a single RARE in the 5’-noncoding region close to exon 1 that is overlapped by or close to the edge of RA-regulated H3K27ac and H3K27me3 peaks (Fig. 4A-B). Recent studies in zebrafish identified RAREs in similar locations in the *nr2f1a* and *nr2f2* genes (Dohn et al. 2019) and this conservation to mouse was detected by our analysis (Table 2).

**Fig. 4.**
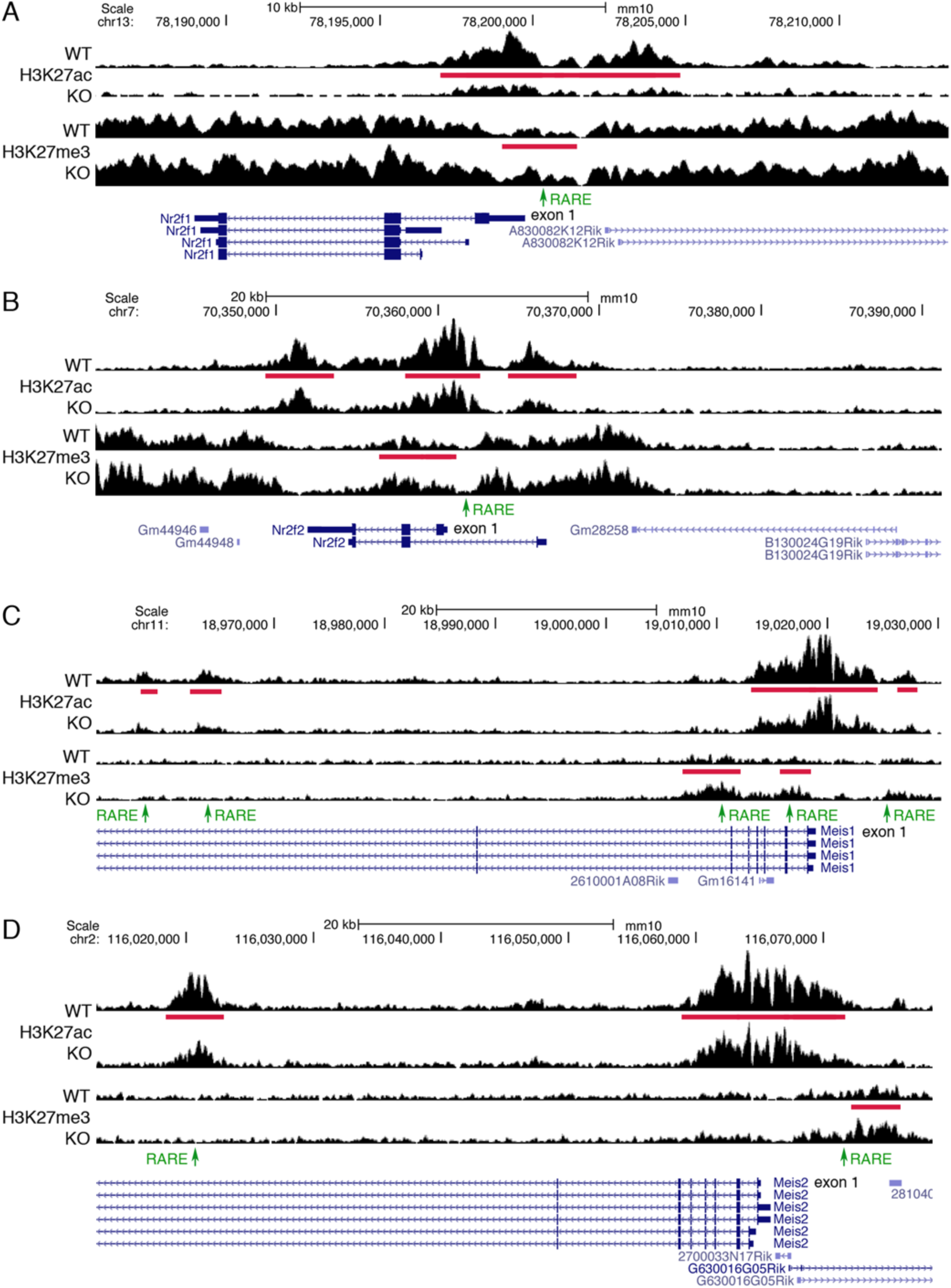
ChIP-seq findings for *Nr2f1*, *Nr2f2*, *Meis1*, and *Meis2* identify RARE enhancers in gene families. (A-B) *Nr2f1* and *Nr2f2* have differential peaks (red bars) for both H3K27ac (decreased) and H3K27me3 (increased) when RA is lost in E8.5 trunk from *Aldh1a2*-/-(KO) compared to wild-type (WT). Each family member has one RARE (green) contained within these differential peaks that are candidates for RARE enhancers. (C-D) *Meis1* and *Meis2* have differential peaks for both H3K27ac (all decreased) and H3K27me3 (all increased) when RA is lost, along with associated RAREs for each peak that are candidates for RARE enhancers.

*Meis1* and *Meis2* were previously shown to be upregulated by RA in chick limbs treated with RA (Mercader et al. 2000), and loss of RA in *Aldh1a2*-/-embryos results in reduced expression of *Meis2* in paraxial mesoderm (Niederreither et al. 2000). Other studies have shown that *Meis1* and *Meis2* are activated by RA in embryonic stem cells and other cell lines, and RAREs were identified in their 5’-noncoding regions (Lalevee et al. 2011; Kashyap et al. 2013). Here, *Meis1* was found to have four RAREs in introns 1, 6 and 7 that are overlapped by RA-regulated peaks for H3K27ac and/or H3K27me3, plus we identified the previously reported RARE in the 5’-noncoding region that is located at the edge of a small RA-regulated H3K27ac peak (Fig. 4C). *Meis2* was found to have two RAREs that are overlapped by RA-regulated peaks for H3K27ac and/or H3K27me3, one in the 5’-noncoding region (previously identified) and another in intron 7 (Fig. 4D). Our analysis shows that *Meis1* and *Meis2* each have a highly conserved DR5 RARE enhancer (Table 2). Together, these studies identify *Nf2f1*, *Nr2f2*, *Meis1*, and *Meis2* as candidate RA target genes in the developing trunk.

### *Nr2f1* and *Nr2f2* function redundantly to control body axis formation

In order to be a biologically important RA target gene, the gene must not only be associated with a RARE, but must perform a function downstream of RA during trunk development which can be determined by gene knockout studies. Here, we sought to validate our approach by performing knockout studies on some of the candidate RA target genes, particularly those that have nearby highly conserved RAREs. One could also undertake deletion studies of the RAREs, but this is only relevant after a knockout of the gene itself shows a defect. In addition, as genes are often controlled by redundant enhancers (which we observed here for many genes that have two or more RAREs associated with RA-regulated epigenetic marks; Table S3), studies in which predicted enhancers are deleted often have no effect on development (Will et al. 2017; Cunningham et al. 2018; Dickel et al. 2018; Osterwalder et al. 2018; Duester 2019); this includes knockout studies we performed on a RARE that was predicted by enhancer transgene studies to be needed for *Tbx5* expression in forelimb bud that had no effect on *Tbx5* or development (Cunningham et al. 2018). Below, we describe gene knockout studies on candidate RA target genes with nearby highly conserved RAREs to determine if these genes have a required function in trunk development.

*Nr2f1* and *Nr2f2* were selected for gene knockout as they both have nearby candidate RARE enhancers (identified by our H3K27ac/H3K27me3 ChIP-seq analysis) that are conserved from mouse to zebrafish (Table 2). *Nr2f1* (formerly known as COUP-TFI) and *Nr2f2* (formerly known as COUP-TFII) are both expressed at E8.5 in somites and presomitic mesoderm but not spinal cord, suggesting they may function in mesoderm formation during body axis formation (Béland and Lohnes 2005; Vilhais-Neto et al. 2010). Here, in situ hybridization analysis shows that *Nr2f1* and *Nr2f2* have reduced expression in the trunk of E8.5 *Aldh1a2*-/-embryos compared to wild-type (Fig. S3).

The *Nr2f1* knockout is lethal at birth with brain defects but no somite, spinal cord, or body axis defects are observed (Qiu et al. 1997). The *Nr2f2* knockout is lethal at E10.5 with defects in heart development but not body axis formation (Pereira et al. 1999). As redundancy may have masked a body axis defect, we generated *Nr2f1/Nr2f2* double mutants. As it would be quite time-consuming and expensive to obtain (if possible) the previously described *Nr2f1* and *Nr2f2* single knockout mouse lines, then generate a double heterozygote mouse line, and then generate double homozygote embryos at a ratio of 1:16, we employed CRISPR/Cas9 gene editing to examine F0 embryos as we previously described for *Ncor1/Ncor2* double mutants (Kumar et al. 2016). Fertilized mouse oocytes were injected with sgRNAs designed to generate frameshift knockout deletions in the second exons of both *Nr2f1* and *Nr2f2*. After dissecting embryos at E9.0, we obtained *Nr2f1/Nr2f2* double knockouts that exhibited a body axis growth defect, more similar in size to that of wild-type E8.25 embryos (Fig. 5). Genotyping showed that embryos carrying 1 or 2 knockout alleles were normal in size compared to E9.0 wild-type (Fig. 5A), whereas embryos carrying either 3 or 4 knockout alleles exhibited a defect in body axis extension and are similar in size to E8.25 wild-type; *n*=7 (Fig. 5B-C). Staining for *Uncx* somite expression demonstrated that embryos with 1-2 knockout alleles all have a normal number of somites with normal size (Fig. 5A), whereas embryos with 3-4 knockout alleles all have less somites that are smaller in size; embryos with 3 knockout alleles (either *Nr2f1*-het/*Nr2f2*-hom or *Nr2f1*-hom/*Nr2f2*-het) or 4 knockout alleles (*Nr2f1*-hom/*Nr2f2*-hom) have a similar small somite defect (Fig. 5B-C). As E9.0 *Nr2f1/Nr2f2* mutants carrying 3-4 knockout alleles are more similar in size to E8.25 wild-type, in order to estimate somite size along the anteroposterior axis we compared them to *Uncx*-stained E8.25 wild-type embryos (Fig. 5D), thus revealing that the E9.0 mutants have somites about 57% the size of somites in E8.25 wild-type embryos, showing they have a specific defect in trunk development rather than a global body growth defect (Fig. 5E).

**Fig. 5.**
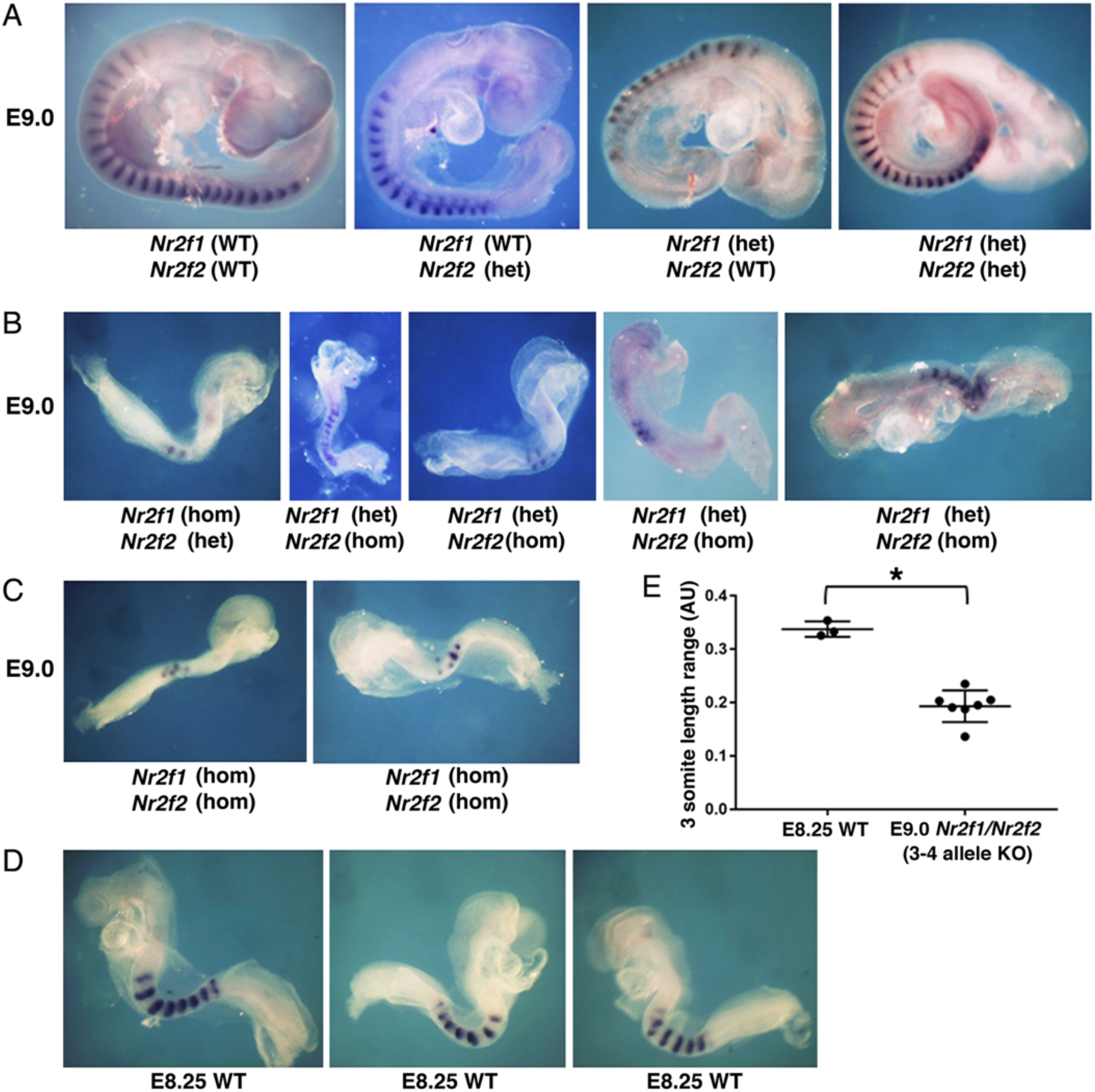
*Nr2f1/Nr2f2* double mutants exhibit defects in body axis formation. (A) Embryos dissected at E9.0 carrying 0-2 knockout alleles for *Nr2f1* or *Nr2f2* have normal somites and body axis formation based on expression of the somite marker *Uncx*. (B-C) Embryos dissected at E9.0 and stained for *Uncx* that carry 3 or 4 knockout alleles for *Nr2f1* or *Nr2f2* exhibit small somites and reduced body axis growth resembling the size of embryos at E8.25. (D) Wild-type (WT) E8.25 embryos stained for *Uncx* expression. (E) Comparison of somite size along the anteroposterior axis between E8.25 WT and E9.0 *Nr2f1/Nr2f2* knockout embryos (3-4 knockout alleles); *, p < 0.05, data expressed as mean ± SD, one-way ANOVA (non-parametric test); WT, *n* = 3 biological replicates; *Nr2f1/Nr2f2* 3-4 allele double knockout, *n* = 7 biological replicates.

Overall, our findings show that loss of 3 or 4 alleles of *Nr2f1* and *Nr2f2* hinders body axis formation and results in smaller somites, thus validating our approach for finding new genes required for body axis formation. Although the body axis defect we observe for *Nr2f1/Nr2f2* double mutants may be caused at least in part by a cardiac defect (Pereira et al. 1999), our RNA-seq and ChIP-seq studies were performed on trunk tissue in which the heart was excluded showing that RA-regulated epigenetic marks are observed near *Nr2f1/Nr2f1* genes located outside the heart in the trunk. Also, our results show that double mutants have smaller somites than wild-type embryos of a comparable size, revealing a specific effect on somitogenesis (body axis extension) rather than just an overall effect on embryonic growth. In the future, more detailed studies of *Nr2f1/Nr2f2* double mutants can be performed to determine how these genes control body axis extension. In addition, future studies can be performed to determine how RARE enhancers function along with other factors to control *Nr2f1* and *Nr2f2* expression during body axis formation.

### *Meis1* and *Meis2* function redundantly to control both body axis and limb formation

*Meis1* and *Meis2* were selected for gene knockout as *Meis1* has a nearby candidate RARE enhancer conserved from mouse to frog, and *Meis2* has a nearby candidate RARE enhancer conserved from mouse to bird (Table 2). *Meis1* and *Meis2* are both expressed throughout the trunk and in the proximal regions of limb buds (Mercader et al. 2000). Here, in situ hybridization analysis shows that *Meis1* and *Meis2* have reduced expression in the trunk of E8.5 *Aldh1a2*-/-embryos compared to wild-type (Fig. S3).

The *Meis1* knockout is lethal at E11.5 with hematopoietic defects, but no body axis or limb defects are observed (Hisa et al. 2004). The *Meis2* knockout is lethal at E14.5 with defects in cranial and cardiac neural crest, but no defects in body axis or limb formation were observed (Machon et al. 2015). As redundancy may have masked a body axis or limb defect, we generated *Meis1/Meis2* double mutants via CRISPR/Cas9 gene editing of fertilized mouse oocytes employing sgRNAs designed to generate frameshift knockout deletions in the second exons of both *Meis1* and *Meis2*. Embryos were dissected at E10.5 and stained for somite *Uncx* expression. Genotyping showed that E10.5 embryos carrying 1 or 2 knockout alleles for *Meis1/Meis2* were normal in size with normal size somites compared to E10.5 wild-type (Fig. 6A). However, E10.5 embryos carrying 3 or 4 knockout alleles for *Meis1/Meis2* exhibited a body axis extension defect and were either similar in size to *Uncx*-stained E9.5 wild-type embryos (*n*=3) or smaller (*n*=4); comparison of somite size along the anteroposterior axis for five of these E10.5 mutants shows that somite sizes range from that seen in E9.5 wild-type to about 40% smaller (Fig. 6B-D). We observed that E10.5 *Meis1/Meis2* mutants carrying 3-4 knockout alleles that grew similar in size and somite number to E9.5 embryos exhibited a lack of forelimb bud outgrowth; *n*=3 (Fig. 6E).

**Fig. 6.**
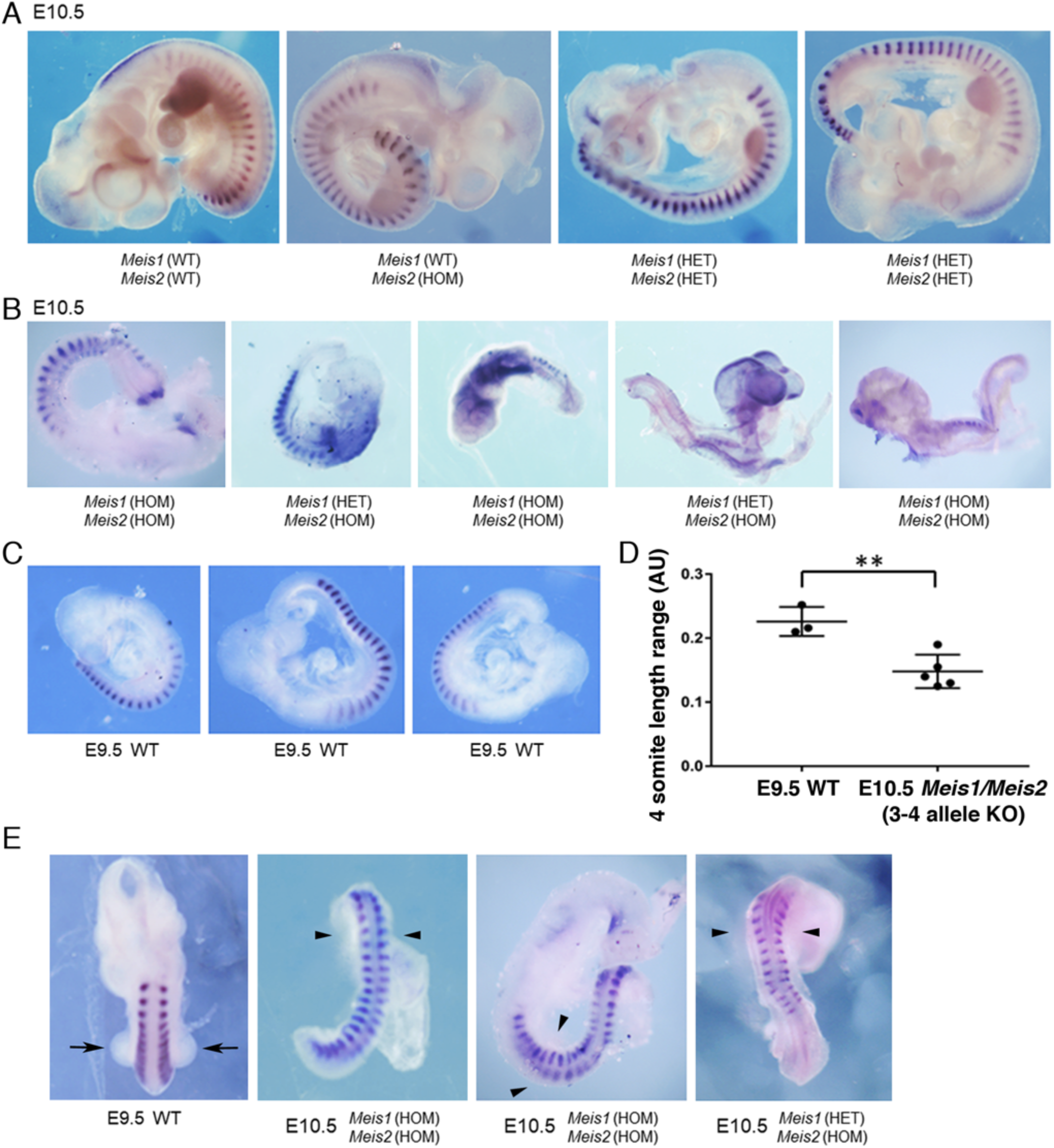
*Mei s1/Meis2* double mutants exhibit defects in body axis and forelimb formation. (A) Embryos dissected at E10.5 carrying 0-2 knockout alleles for *Meis1* or *Meis2* have normal somites and body axis formation based on expression of the somite marker *Uncx;* also, limb formation is normal. (B) Embryos dissected at E10.5 and stained for *Uncx* that carry 3 or 4 knockout alleles for *Meis1* or *Meis2* exhibit small somites and reduced body axis growth resembling the size of embryos at E9.5. (C) Wild-type (WT) E9.5 embryos stained for *Uncx* expression. (D) Comparison of somite size along the anteroposterior axis between E9.5 WT and E10.5 *Meis1/Meis2* knockout embryos (3-4 knockout alleles); *, p < 0.05, data expressed as mean ± SD, one-way ANOVA (non-parametric test); WT, *n* = 3 biological replicates; *Meis1/Meis2* 3-4 allele double knockout, *n* = 5 biological replicates. (E) Forelimb buds (arrows) normally observed in an E9.5 WT embryo are absent in E10.5 *Meis1/Meis2* knockout embryos with 3-4 knockout alleles (arrowheads}.

Overall, our findings show that loss of 3 or 4 alleles of *Meis1* and *Meis2* hinders body axis and forelimb formation, thus providing further evidence that our method of identifying candidate RARE enhancers can identify genes essential for development. In the future, more detailed studies of *Meis1/Meis2* double mutants can be performed to determine how these genes control body axis and limb formation, plus additional studies can be performed to determine how RARE enhancers function along with other factors to control *Meis1* and *Meis2* expression during early development.

## Discussion

Our epigenetic ChIP-seq studies combined with RNA-seq on wild-type vs *Aldh1a2*-/-RA-deficient trunk tissue provides a means for identifying new candidate RA target genes that may be required for development. By focusing on RA-regulated genes that also have changes in nearby RA-regulated H3K27ac and/or H3K27me3 epigenetic marks associated with highly conserved RARE enhancers or silencers, our approach can be used to identify excellent candidates for gene knockout studies to learn more about gene function.

Here, in our studies on *Aldh1a2*-/-trunk tissue, we were able to narrow down 4298 genes identified with RNA-seq that have significant changes in gene expression following loss of RA to 38 excellent candidate RA target genes in E8.5 trunk that also have significant changes in H3K27ac and/or H3K27me3 marks (located nearby or further away in the same TAD) associated with highly conserved RAREs. Our method allows one to identify genes that are most likely to be transcriptional targets of the RA signaling pathway as opposed to those whose expression is changed by effects downstream of RARs and RA signaling such as changes in expression or activity of other transcription factors or post-transcriptional changes in mRNA abundance. Our findings allow us to predict that some genes are likely to be indirect transcriptional targets of RA as they have nearby RA-regulated peaks for H3K27ac or H3K27me3 but no RAREs, i.e. *Pax6* that is transcriptionally regulated by factors whose expression is altered by loss of RA including *Sox2* (Oosterveen et al. 2013), *Cdx* (Joshi et al. 2019), and *Fgf8* (Patel et al. 2013).

Our findings provide evidence for additional RARE silencers. Previous methods designed to identify RAREs favored discovery of RARE enhancers as studies were designed to find DNA elements that when fused to a heterologous promoter and marker gene would stimulate expression of the marker gene in the presence of RA. Also, when nuclear receptor coactivators (NCOA) and corepressors (NCOR) that control RA signaling were originally discovered, the model proposed for their function suggested that binding of RA to RAR favored binding of NCOA to activate transcription, with unliganded RAR favoring release of NCOA and binding of NCOR to repress transcription (Perissi et al. 2004). However, analysis of the *Fgf8* RARE silencer at −4.1 kb demonstrated that RARs bound to RAREs can recruit NCOR in an RA-dependent manner, plus this RARE is required for normal body axis extension (Kumar et al. 2016). The *Fgf8* RARE silencer was also found to recruit Polycomb Repressive Complex 2 (PRC2) and histone deacetylase 1 (HDAC1) in an RA-dependent manner, providing further evidence that RA can directly control gene silencing (Kumar and Duester 2014). Here, we identified additional RARE silencers near *Fgf8* and *Cdx2* plus several additional genes. Our studies indicate that RARE silencers are less common than RARE enhancers, and we found that *Fgf8* is the only gene associated with a RARE silencer conserved beyond mammals. These additional RARE silencers can be further examined in comparison to the *Fgf8* RARE silencer to determine the mechanism through which RA directly represses transcription. It will be important to determine how RAREs can function as RA-dependent enhancers for some genes but RA-dependent silencers for other genes.

RA has been shown to be required for balanced NMP differentiation during body axis formation by favoring a neural fate over a mesodermal fate (Cunningham et al. 2015; Henrique et al. 2015; Gouti et al. 2017). Our studies provide evidence that RA directly regulates several genes at the trunk/caudal border needed for NMP differentiation; i.e. activation of *Sox2* in the neural plate that favors neural differentiation, repression of *Fgf8* that favors mesodermal differentiation, and repression of *Cdx2* that helps define the location of NMPs. We now provide evidence for a candidate RARE enhancer that activates *Sox2*, three candidate RARE silencers that repress *Cdx2*, and two additional candidate RARE silencers for *Fgf8*. As the knockout of the original *Fgf8* RARE silencer at −4.1 kb exhibited a body axis phenotype less severe than loss of RA in *Aldh1a2*-/-embryos (Kumar et al. 2016), it is possible that the additional two candidate RARE silencers found here provide redundant functions for *Fgf8* repression.

Our observation of highly conserved candidate RARE enhancers near two members of two different gene families (*Nr2f* and *Meis*) was intriguing as it suggested that these gene family members may play redundant roles in body axis formation downstream of RA. As we were not sure whether the lack of previous studies on *Nf2f1;Nf2f2* double knockout and *Meis1;Meis2* double knockout mouse lines may be due to defects in double heterozygote adults that prevent generation of double homozygote embryos by conventional genetic approaches, we employed CRISPR gene editing to directly generate F0 double knockouts. Our *Nr2f1/Nr2f2* double knockout studies indeed revealed a defect in body axis formation and small somites that is not observed in each single knockout. Interestingly, zebrafish *nr2f1a/nr2f2* double knockout embryos reported recently exhibit a heart defect more severe than each single knockout, but not a body axis defect or body growth defect (Dohn et al. 2019). This observation is consistent with studies showing that RA is not required for NMP differentiation or body axis formation in zebrafish (Begemann et al. 2001; Berenguer et al. 2018). Thus, it appears that the ancestral function of *Nr2f* genes in fish was to control heart formation, but that during evolution another function to control body axis formation was added. Future studies can be directed at understanding the mechanism through which *Nr2f1* and *Nr2f2* control body axis formation.

The *Meis1/Meis2* double knockouts we describe here revealed an unexpected function for *Meis* genes in body axis extension and forelimb initiation. *Meis1* and *Meis2* are markers of the proximal limb during forelimb and hindlimb development and were proposed to be activated by RA in the proximal limb as part of the proximodistal limb patterning mechanism in chick embryos (Mercader et al. 2000; Cooper et al. 2011; Rosello-Diez et al. 2011). However, knockout of *Rdh10* required to generate RA demonstrated that complete loss of RA in the limb fields prior to and during limb development did not affect hindlimb initiation or patterning, whereas forelimbs were stunted but with *Meis1* and *Meis2* expression still maintained in a proximal position in both stunted forelimbs and hindlimbs (Cunningham et al. 2011; Cunningham et al. 2013); reviewed in (Cunningham and Duester 2015). Our epigenetic results here support the previous proposal that RA can up-regulate *Meis1* and *Meis2* (but in the body axis prior to limb formation as opposed to the limb itself) and we provide evidence that *Meis1* and *Meis2* are transcriptional targets of RA in the body axis. Future studies can be directed at understanding the mechanism through which *Meis1* and *Meis2* control body axis and limb formation.

Our studies demonstrate the power of combining gene knockouts, ChIP-seq on epigenetic marks, and RNA-seq to identify transcription factor target genes required for a particular developmental process. In addition to H3K27ac and H3K27me3 epigenetic marks that are quite commonly observed near genes during activation or repression, respectively, it is likely that further ChIP-seq studies that identify RA-regulated binding sites for coactivators and corepressors will provide additional insight into RA target genes and transcriptional pathways. Such knowledge is essential for determining the mechanisms through which RA controls developmental pathways and should be useful to address RA function in adult organs. A similar epigenetic approach can be used to determine the target genes for any transcriptional regulator for which a knockout is available, thus accelerating the ability to understand gene regulatory networks in general.

## Methods

### Generation of *Aldh1a2*-/-mouse embryos and isolation of trunk tissue

*Aldh1a2*-/-mice have been previously described (Mic et al. 2002). E8.5 *Aldh1a2*-/-embryos were generated via timed matings of heterozygous parents; genotyping was performed by PCR analysis of yolk sac DNA. E8.5 trunk tissue was released from the rest of the embryo by dissecting across the posterior hindbrain (to remove the head, anterior hindbrain, pharyngeal region, and heart) and just posterior to the most recently formed somite (to remove the caudal progenitor zone) as previously described (Kumar and Duester 2014). All mouse studies conformed to the regulatory standards adopted by the Institutional Animal Care and Use Committee at the SBP Medical Discovery Institute which approved this study under Animal Welfare Assurance Number A3053-01 (approval #18-092).

### RNA-seq analysis

Total RNA was extracted from E8.5 trunk tissue (two wild-type trunks and two *Aldh1a2*-/-trunks) and DNA sequencing libraries were prepared using the SMARTer Stranded Total RNA-Seq Kit v2 Pico Input Mammalian (Takara). Sequencing was performed on Illumina NextSeq 500, generating 40 million reads per sample with single read lengths of 75 bp. Sequences were aligned to the mouse mm10 reference genome using TopHat splice-aware aligner; transcript abundance was calculated using Expectation-Maximization approach; fragments per kilobase of transcript per million mapped reads (FPKM) was used for sample normalization; Generalized Linear Model likelihood ratio test in edgeR software was used as a differential test. High throughput DNA sequencing was performed in the Sanford Burnham Prebys Genomics Core.

### qRT-PCR analysis

Total RNA was extracted from 20 trunks of either E8.5 wild-type or *Aldh1a2*-/-embryos with the RNeasy Micro Kit (Qiagen #74004). Reverse transcription was performed with the High-Capacity cDNA RT Kit (Thermo Fisher Scientific #4368814). Quantitative PCR (qPCR) was performed using Power SYBR Green PCR Master Mix (Life Tech Supply #4367659). Relative quantitation was performed using the ddCt method with the control being expression of *Rpl10a*. Primers used for PCR (5’-3’):

Rpl10a-F ACCAGCAGCACTGTGATGAA

Rpl10a-R cAGGATACGTGGgATCTGCT

Rarb-F CTCTCAAAGCCTGCCTCAGT

Rarb-R GTGGTAGCCCGATGACTTGT

Nr2f1-F TCAGGAACAGGTGGAGAAGC

Nr2f1-R ACATACTCCTCCAGGGCACA

Nr2f2-F GACTCCGCCGAGTATAGCTG

Nr2f2-R GAAGCAAGAGCTTTCCGAAC

Meis1-F CAGAAAAAGCAGTTGGCACA

Meis1-R TGCTGACCGTCCATTACAAA

Meis2-F AACAGTTAGCGCAAGACACG

Meis2-R GGGCTGACCCTCTGGACTAT

Spry4-F CCTGTCTGCTGTGCTACCTG

Spry4-R AAGGCTTGTCAGACCTGCTG

### Chromatin immunoprecipitation (ChIP) sample preparation for ChIP-seq

For ChIP-seq we used trunk tissue from E8.5 wild-type or *Aldh1a2*-/-embryos dissected in modified PBS, i.e. phosphate-buffered saline containing 1X complete protease inhibitors (concentration recommended by use of soluble EDTA-free tablets sold by Roche #11873580001) and 10 mM sodium butyrate as a histone deacetylase inhibitor (Sigma # B5887). Samples were processed similar to previous methods (Voss et al. 2012). Dissected trunks were briefly centrifuged in 1.5 ml tubes and excess PBS dissection buffer was removed. For cross-linking of chromatin DNA and proteins, 500 μl 1% formaldehyde was added, the trunk samples were minced by pipetting up and down with a 200 μl pipette tip and then incubated at room temperature for 15 min. To stop the cross-linking reaction, 55 μl of 1.25 M glycine was added and samples were rocked at room temperature for 5 min. Samples were centrifuged at 5000 rpm for 5 min and the supernatant was carefully removed and discarded. A wash was performed in which 1000 μl of ice-cold modified PBS was added and mixed by vortex followed by centrifugation at 5000 rpm for 5 min and careful removal of supernatant that was discarded. This wash was repeated. Cross-linked trunk samples were stored at −80C until enough were collected to proceed, i.e. 100 wild-type trunks and 100 *Aldh1a2*-/-trunks to perform ChIP-seq with two antibodies in duplicate.

Chromatin was fragmented by sonication. Cross-linked trunk samples were pooled, briefly centrifuged, and excess PBS removed. 490 μl lysis buffer (modified PBS containing 1% SDS, 10 mM EDTA, 50 mM Tris-HCl, pH 8.0) was added, mixed by vortexing, then samples were incubated on ice for 10 min. Samples were divided into four sonication microtubes (Covaris AFA Fiber Pre-Slit Snap-Cap 6×16 mm, #520045) with 120 μl per tube. Sonication was performed with a Covaris Sonicator with the following settings - Duty: 5%, Cycle: 200, Intensity: 4, #Cycles: 10, 60 sec each for a total sonication time of 14 min. The contents of the four tubes were re-combined by transfer to a single 1.5 ml microtube which was then centrifuged for 10 min at 13,000 rpm and the supernatants transferred to a fresh 1.5 ml microtube. These conditions were found to shear trunk DNA to an average size of 300 bp using a 5 μl sample for Bioanalyzer analysis. At this point 20 μl was removed for each sample (wild-type trunks and *Aldh1a2*-/-trunks) and stored at −20C to serve as input DNA for ChIP-seq.

Each sample was divided into four 100 μl aliquots to perform immunoprecipitation with two antibodies in duplicate. Immunoprecipitation was performed using the Pierce Magnetic ChIP Kit (Thermo Scientific, #26157) following the manufacturer’s instructions and ChIP-grade antibodies for H3K27ac (Active Motif, Cat#39133) or H3K27me3 (Active Motif, Cat#39155). The immunoprecipitated samples and input samples were subjected to reversal of cross-linking by adding water to 500 μl and 20 μl 5 M NaCl, vortexing and incubation at 65C for 4 hr; then addition of 2.6 μl RNase (10 mg/ml), vortexing and incubation at 37C for 30 min; then addition of 10 μl 0.5 M

EDTA, 20 μl 1 M Tris-HCl, pH 8.0, 2 μl proteinase K (10 mg/ml), vortexing and incubation at 45C for 1 hr. DNA was extracted using ChIP DNA Clean & Concentrator (Zymo, # D5201 & D5205), After elution from the column in 50 μl of elution buffer, the DNA concentration was determine using 2 μl samples for Bioanalyzer analysis. The two input samples ranged from 16-20 ng/μl and the eight immunoprecipitated samples ranged from 0.1-0.2ng/μl (5-10 ng per 100 trunks). For ChIP-seq, 2 ng was used per sample to prepare libraries for DNA sequencing.

### ChIP-seq genomic sequencing and bioinformatic analysis

Libraries for DNA sequencing were prepared according to the instructions accompanying the NEBNext DNA Ultra II kit (catalog # E7645S; New England Biolabs, Inc). Libraries were sequenced on the NextSeq 500 following the manufacturer’s protocols, generating 40 million reads per sample with single read lengths of 75 bp. Adapter remnants of sequencing reads were removed using cutadapt v1.18 (Martin 2011). ChIP-Seq sequencing reads were aligned using STAR aligner version 2.7 to Mouse genome version 38 (Dobin et al. 2013). Homer v4.10 (Heinz et al. 2010) was used to call peaks from ChIP-Seq samples by comparing the ChIP samples with matching input samples. Homer v4.10 was used to annotate peaks to mouse genes, and quantify reads count to peaks. The raw reads count for different peaks were compared using DESeq2 (Love et al. 2014). P values from DESeq2 were corrected using the Benjamini & Hochberg (BH) method for multiple testing errors (Benjamini and Hochberg 1995). Peaks with BH corrected p value <0.05 (BHP<0.05) were selected as significantly differentially marked peaks. Transcription factor binding sites motif enrichment analyses were performed using Homer v4.10 (Heinz et al. 2010) to analyze the significant RA-regulated ChIP-seq peaks; DR1 RAREs were found by searching for TR4(NR),DR1; DR2 RAREs by Reverb(NR),DR2; and DR5 RAREs by RAR:RXR(NR),DR5. Evolutionary conservation of RAREs was performed via DNA sequence homology searches using the UCSC genome browser software. TAD analysis was performed using the 3D Genome Browser (http://promoter.bx.psu.edu/hi-c/view.php); we used the TAD database from Hi-C data for mouse ES cells reported for the mouse mm10 genome. Ingenuity Pathway Analysis (IPA) was used to identify pathways for our list of target genes; from IPA results, heatmaps were designed with Prism software and associated networks were created using STRING software. High throughput DNA sequencing was performed in the Sanford Burnham Prebys Genomics Core and bioinformatics analysis was performed in the Sanford Burnham Prebys Bioinformatics Core.

### Generation of mutant embryos by CRISPR/Cas9 mutagenesis

CRISPR/Cas9 gene editing was performed using methods similar to those previously described by others (Wang et al. 2013; Tan et al. 2015) and by our laboratory (Kumar et al. 2016). Single-guide RNAs (sgRNAs) were generated that target exons to generate frameshift null mutations, with two sgRNAs used together for each gene. sgRNAs were designed with maximum specificity using the tool at crispr.mit.edu to ensure that each sgRNA had no more than 17 out of 20 matches with any other site in the mouse genome and that those sites are not located within exons of other genes. DNA templates for sgRNAs were generated by PCR amplification (Phusion DNA Polymerase; New England Biolabs) of ssDNA oligonucleotides (purchased from Integrated DNA Technologies) containing on the 5’ end a minimal T7 promoter, then a 20 nucleotide sgRNA target sequence (underlined below), and finally the tracrRNA sequence utilized by Cas9 on the 3’ end, shown as follows:

5’-GCGTAATACGACTCACTATAGG<u>NNNNNNNNNNNNNNNNNNNN</u>GTTTTAGAGCTAGAAATA GCAAGTTAAAATAAGGCTAGTCCGTTATCAACTTGAAAAAGTGGCACCGAGTCGGTGCTTTT-3’

The 20 nucleotide target sequences used were as follows:

*Nf2f1* exon 2 (#1) TTTTTATCAGCGGTTCAGCG

*Nf2f1* exon 2 (#2) GGTCCATGAAGGCCACGACG

*Nf2f2* exon 2 (#1) GGTACGAGTGGCAGTTGAGG

*Nf2f2* exon 2 (#2) CGCCGAGTATAGCTGCCTCA

*Meis1* exon 2 (#1) CGACGACCTACCCCATTATG

*Meis1* exon 2 (#2) TGACCGAGGAACCCATGCTG

*Meis2* exon 2 (#1) GATGAGCTGCCCCATTACGG

*Meis2* exon 2 (#2) CGACGCCTTGAAAAGAGACA

sgRNAs were then transcribed from templates using HiScribe T7 High Yield RNA Synthesis Kit (New England Biolabs) and purified using Megaclear Kit (Life Technologies). sgRNAs were tested in vitro for their cleavage ability in combination with Cas9 nuclease (New England Biolabs); briefly, genomic regions flanking the target sites were PCR amplified, then 100 ng was incubated with 30 nM Cas9 nuclease and 30 ng sgRNA in 30 µl for 1 hour at 37°C, followed by analysis for cleavage by gel electrophoresis.

For injection into mouse embryos, a solution containing 50 ng/µl Cas9 mRNA (Life Technologies) and 20 ng/µl for each sgRNA used was prepared in nuclease free water. Fertilized oocytes were collected from 3-4 week-old superovulated C57Bl6 females prepared by injecting 5 IU each of pregnant mare serum gonadotrophin (PMSG) (Sigma Aldrich) and human chorionic gonadotropin (hCG) (Sigma Aldrich). Fertilized oocytes were then transferred into M2 medium (Millipore) and injected with the Cas9 mRNA/sgRNA solution into the cytoplasm. Injected embryos were cultured in KSOMaa medium (Zenith) in a humidified atmosphere with 5% CO_2_ at 37°C overnight to maximize the time for CRISPR/Cas9 gene editing to occur at the 1-cell stage, then re-implanted at the 2-cell stage into recipient pseudo-pregnant ICR female mice. Implanted females were sacrificed to obtain F0 E9.0 embryos (*Nr2f1/Nr2f2*) or F0 E10.5 embryos (*Meis1/Meis2*). As fertilized mouse oocytes spend a long time at the 1-cell and 2-cell stages, this facilitates CRISPR/Cas9 gene editing at early stages and allows many F0 embryos to be examined for mutant phenotypes (Kumar et al. 2016). For genotyping, yolk sac DNA was collected and PCR products were generated using primers flanking the sgRNA target sites; PCR products were subjected to DNA sequence analysis from both directions using either upstream or downstream primers. For each gene analyzed, embryos were classified as heterozygous (het) if the DNA sequence contained both a wild-type allele and a frame-shift allele; embryos were classified as homozygous (hom) if only frame-shift alleles were detected but no wild-type sequence.

### Body axis length analysis of embryos

ImageJ software (https://imagej.net) (Schneider et al. 2012) was used to measure body axis length along a 3-somite region (*Nr2f1/Nr2f2* double mutants) or 4-somite region (*Meis1/Meis2* double mutants) compared to wild-type, with each specimen photographed at the same magnification. Statistical analysis was performed using one-way ANOVA (non-parametric test) with data presented as mean ± standard deviation (SD) and with p > 0.05 indicating significance.

### In situ gene expression analysis

Embryos were fixed in paraformaldehyde at 4°C overnight, dehydrated into methanol, and stored at −20°C. Detection of mRNA was performed by whole mount in situ hybridization as previously described (Sirbu and Duester 2006).

## Acknowledgments

We thank the Genomics Core Facility and Bioinformatics Core Facility at SBP Medical Discovery Institute for help with ChIP-seq and RNA-seq analysis. We thank the Animal Resources Core Facility at SBP Medical Discovery Institute for conducting timed-matings to generate mouse embryos. Research reported in this publication was supported by NIAMS of the National Institutes of Health under award number R01AR067731 (G.D.). 100%/$2,145,500 of the total project costs were financed with Federal funding. 0%/$0 of the total costs were financed with non-Federal funding. The content is solely the responsibility of the authors and does not necessarily represent the official views of the National Institutes of Health.

## Author Contributions

M.B, K.F.M., and G.D. designed the study and performed the experiments. M.B., J.Y., and G.D. analyzed the data and wrote the paper.

## Data availability

RNA-seq data have been deposited in GEO under accession number GSE131584. ChIP-seq data have been deposited in GEO under accession number GSE131624. All other relevant data are within the paper and its Supporting Information files.

---

**Supplemental Figure S1.**
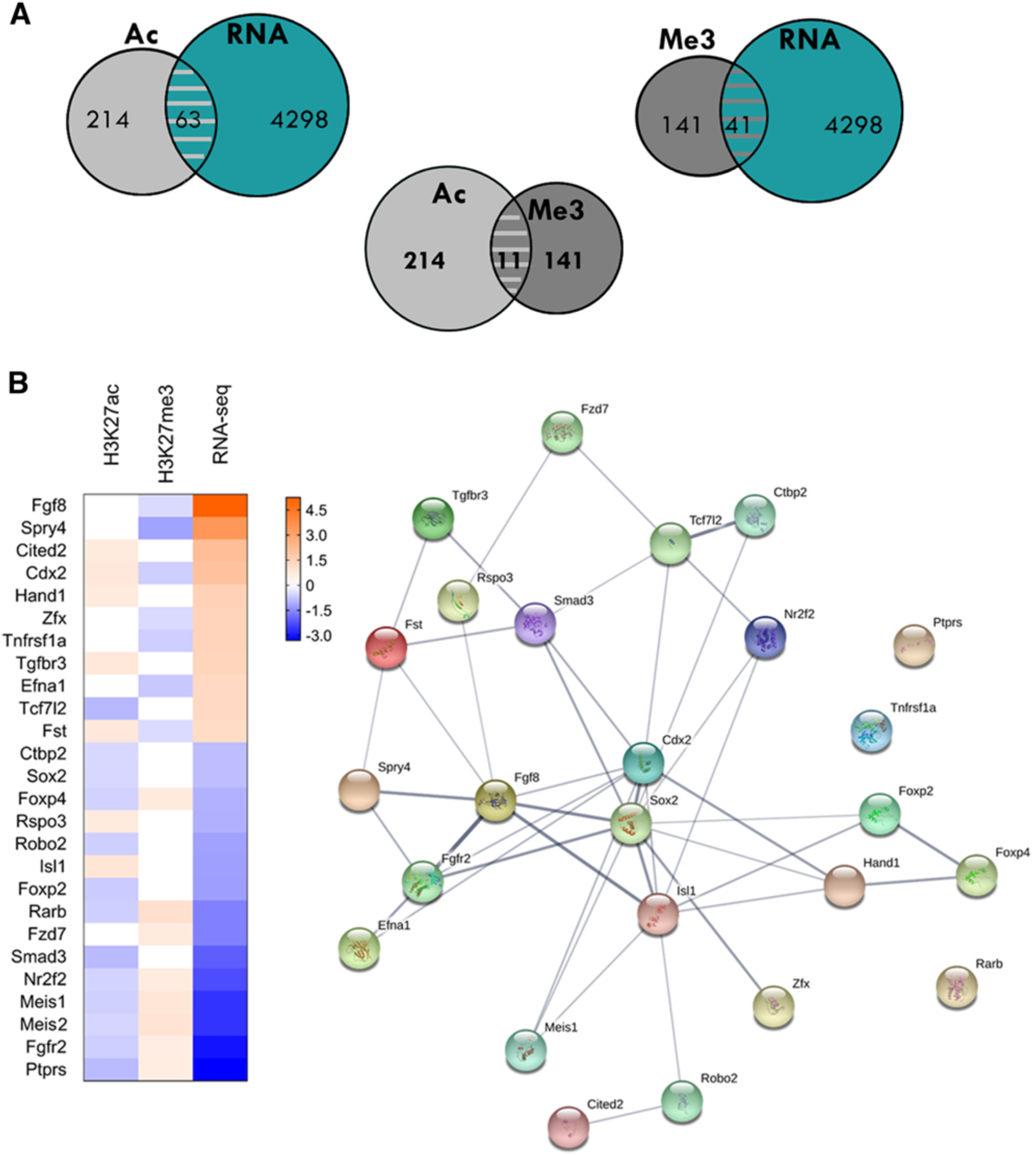
Bioinformatic analysis of genes identified as RA target genes. (A) Venn diagram showing the number of genes that have both RA-regulated expression and RA­ regulated deposition of nearby H3K27a or H3K27me3 marks following loss of RA. (B) The heatmap was designed with Prism software (left panel) from the list of genes involved in “Development of Body Trunk” obtained by IPA analysis of RA target genes identified by loss of RA, and the associated network was created using STRING software (right panel).

**Supplemental Figure 52.**
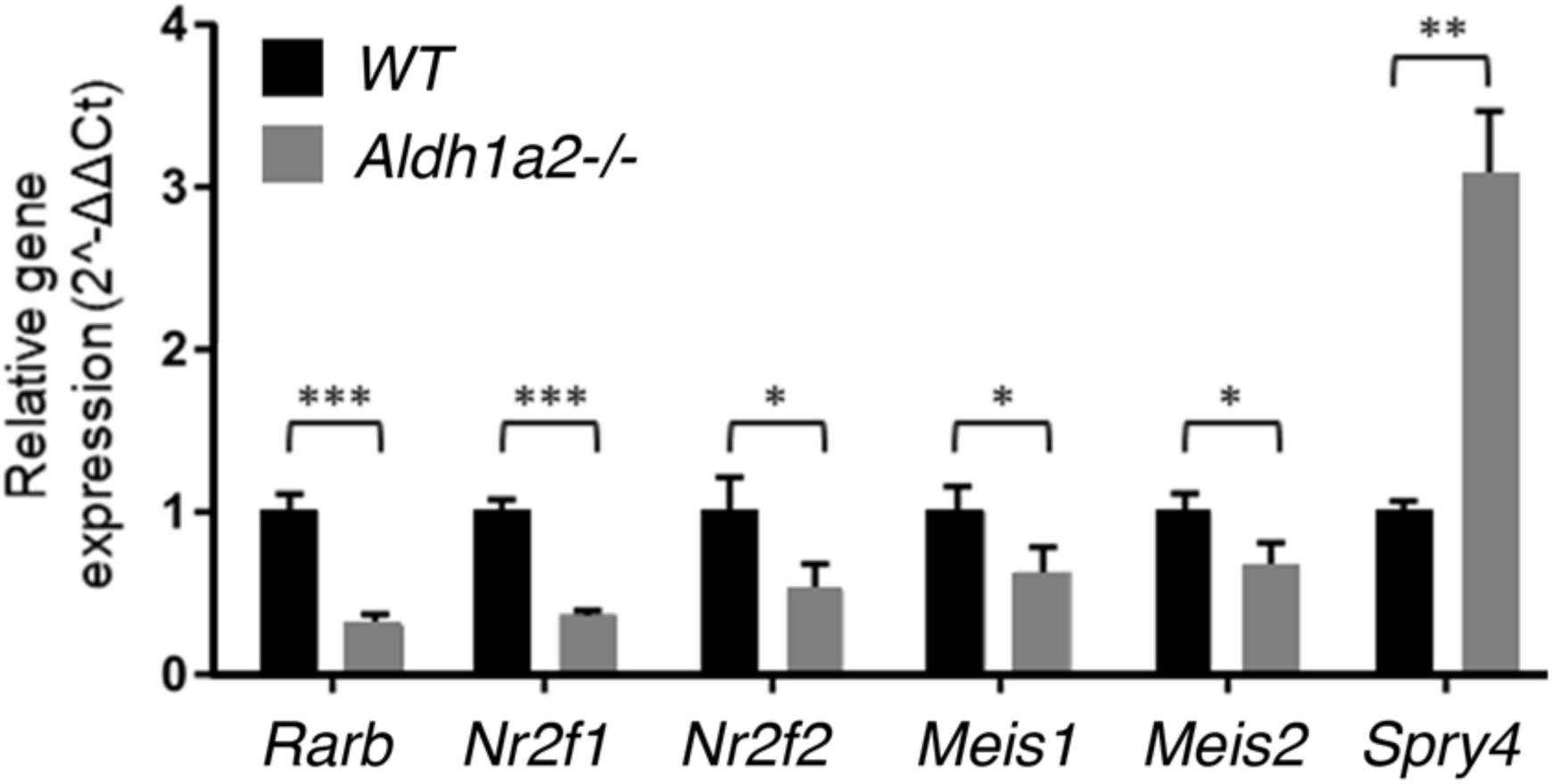
Analysis of differential gene expression of new RA target genes by qRT-PCR analysis of E8.5 wild-type vs *Aldh1a2-l-trunk* tis sue.

**Supplemental Figure 53.**
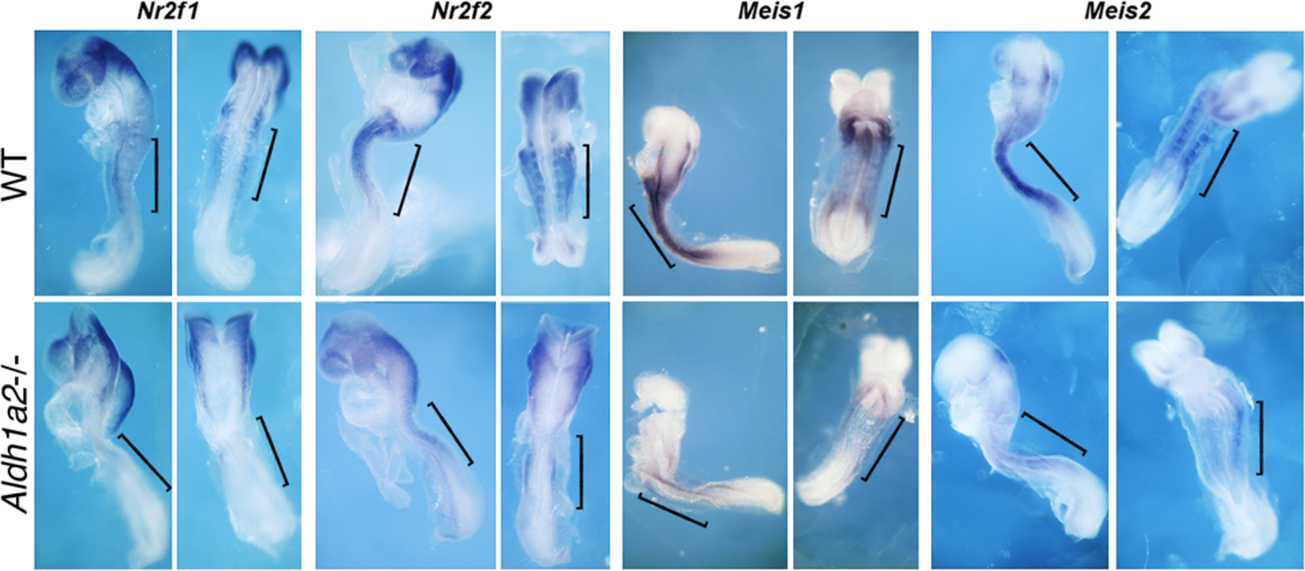
A**n**alysis **of *Nr2f* and *Meis* expression by in situ hybridization.** Analysis of *Nr2f1, Nr2f2, Meis1,* and *Meis2* expression by whole-mount in situ hybridization of E8.5 wild-type (WT} vs *Aldh1a2-/-* trunk tissue; n = 3 for both WT and *Aldh1a2-I-;* brackets point to trunk tissue; for each gene analyzed, both lateral (left) and dorsal (right) views are shown, except ventral for *Meis1*.

**Supplemental Table S1.** Comparison of *Aldh1a2-/-* and wild-type E8.5 trunk tissue for H3K27ac ChIP-seq and RNA-seq results to identify RA-regulated H3K27ac ChIP-seq peaks near genes with RA-regulated expression.

| H3K27ac ChIP-seq differential peak for *Aldh1a2* KO vs WT (mm10) | log2 fold change: H3K27ac ChIP-seq for *Aldh1a2* KO vs WT | RARE: based on Homer TFBS analysis | nearby gene with altered expression in *Aldh1a2* KO vs WT | log2 fold change for nearby gene: RNA-seq for *Aldh1a2* KO vs WT |
| --- | --- | --- | --- | --- |
| chr13:78197222-78204291 | -1.23 | DR1 | Nr2f1 | -2.02 |
| chr13:34133342-34134366 | -1.13 | - | Tubb2b | -1.03 |
| chr18:83838984-83841358 | -1.10 | DR5, DR2 | Tshz1 | -1.33 |
| chr11:68049983-68051745 | -1.07 | DR2, DR1 | Stx8 | -2.87 |
| chr6:84975748-84978033 | -1.03 | DR5 | Zfp638 | -1.33 |
| chr8:14991303-14993068 | -0.96 | - | Arhgef10 | -1.56 |
| chr11:68088685-68091257 | -0.90 | - | Stx8 | -2.87 |
| chr17:56468893-56471601 | -0.90 | DR2, DR1 | Ptprs * | -3.31 |
| chr9:63715651-63717278 | -0.89 | DR5 | Smad3 | -2.09 |
| chr19:21164673-21168435 | -0.86 | DR2 | Zfand5 | -2.37 |
| chr9:16152232-16154557 | -0.86 | - | Fat3 | -3.02 |
| chr9:118653618-118656175 | -0.84 | DR5, DR1 | Itga9 | -1.63 |
| chr6:5028587-5031108 | -0.83 | DR5, DR2 | Ppp1r9a | -1.47 |
| chr9:114798509-114801621 | -0.81 | - | Cmtm8 | -2.12 |
| chr4:149254383-149256741 | -0.80 | - | Kif1b | -1.72 |
| chr5:36373121-36374553 | -0.79 | DR5 | Sorcs2 | -5.20 |
| chr3:5387155-5389128 | -0.79 | DR5 | Zfhx4 * | -2.26 |
| chr17:30467016-30471968 | -0.78 | DR1 | Btbd9 | -1.18 |
| chr3:5235978-5239655 | -0.77 | - | Zfhx4 * | -2.26 |
| chr9:35442832-35445303 | -0.74 | - | Cdon | -1.60 |
| chr14:98034746-98040239 | -0.71 | DR5, DR1 | Dach1 | -2.98 |
| chr6:14897860-14903160 | -0.67 | DR1 | Foxp2 | -1.26 |
| chr16:44529549-44532004 | -0.67 | - | Boc | -0.87 |
| chr14:52330133-52333035 | -0.67 | - | Sall2 | -1.17 |
| chr4:145033496-145035860 | -0.65 | DR5 | Dhrs3 * | -1.11 |
| chr12:8912431-8914944 | -0.64 | - | Laptm4a | -1.97 |
| chr18:83927674-83929824 | -0.64 | - | Tshz1 | -1.33 |
| chr14:14069903-14072065 | -0.64 | - | Atxn7 | -1.50 |
| chr5:111242680-111244777 | -0.63 | DR1 | Ttc28 | -1.87 |
| chr17:66410064-66414806 | -0.63 | DR5, DR1 | Mtcl1 | -1.32 |
| chr4:148106909-148109328 | -0.63 | - | Draxin | -3.43 |
| chr14:16571405-16576397 | -0.63 | DR5 | Rarb * | -1.64 |
| chr16:74395975-74399535 | -0.63 | DR1 | Robo2 | -1.22 |
| chr18:84069541-84075594 | -0.62 | DR5 | Tshz1 | -1.33 |
| chr11:18962656-18965461 | -0.61 | DR5, DR1 | Meis1 * | -2.64 |
| chr9:96989027-96991630 | -0.60 | DR5 | Spsb4 | -2.23 |
| chr3:108409804-108412280 | -0.60 | - | Celsr2 | -7.29 |
| chr17:47897351-47899993 | -0.59 | DR1 | Foxp4 * | -1.02 |
| chr14:78848672-78851835 | -0.58 | DR5, DR2, DR1 | Vwa8 | -1.08 |
| chr2:105689278-105690982 | -0.58 | - | Pax6 | -3.02 |
| chr3:87956774-87961235 | -0.58 | DR2, DR1 | Crabp2 | -2.82 |
| chr2:116019003-116024272 | -0.58 | DR2 | Meis2 * | -1.10 |
| chr4:107834342-107836683 | -0.58 | DR2 | Lrp8 | -1.54 |
| chr7:70348715-70369942 | -0.57 | DR1 | Nr2f2 * | -2.32 |
| chr11:18956989-18958835 | -0.57 | DR5 | Meis1 * | -2.64 |
| chr13:34129519-34132640 | -0.55 | DR2 | Tubb2b | -1.03 |
| chr11:19012000-19025444 | -0.54 | DR1 | Meis1 * | -2.64 |
| chr9:96956410-96959728 | -0.54 | DR2 | Spsb4 | -2.23 |
| chr9:48692264-48699040 | -0.54 | DR2, DR1 | Zbtb16 | 1.36 |
| chr3:34678267-34680699 | -0.54 | DR2 | Sox2 | -0.86 |
| chr6:144250107-144252835 | -0.52 | - | Sox5 | -2.33 |
| chr18:61033064-61036494 | -0.52 | DR2, DR1 | Cdx1 | -2.00 |
| chr3:34647848-34655776 | -0.51 | - | Sox2 | -0.86 |
| chr17:56475307-56476820 | -0.51 | - | Ptprs * | -3.30 |
| chr7:133035031-133040386 | -0.51 | DR2 | Ctbp2 | -3.06 |
| chr18:53463017-53465407 | 0.53 | - | Prdm6 | 0.93 |
| chr19:45733505-45735997 | 0.53 | DR1 | Fgf8 * | 5.24 |
| chr11:54891361-54894784 | 0.57 | DR1 | Gpx3 | 2.58 |
| chr4:86669294-86671201 | 0.59 | - | Plin2 | 2.29 |
| chr3:127457093-127465482 | 0.62 | - | Ank2 | 2.26 |
| chr18:60492434-60494743 | 0.62 | - | Smim3 | 1.25 |
| chr10:17704835-17706602 | 0.65 | DR1 | Cited2 | 2.13 |
| chr11:57831171-57833707 | 0.66 | - | Hand1 | 1.46 |
| chr6:52310576-52314619 | 0.71 | DR1 | Evx1 | 1.54 |
| chr13:114456392-114460659 | 0.72 | DR2 | Fst * | 1.15 |
| chr5:147298587-147311126 | 0.73 | DR2 | Cdx2 * | 1.98 |
| chr5:53106977-53110254 | 0.74 | DR1 | Sel1l3 | 3.67 |
| chr10:59957002-59959223 | 0.75 | - | Ddit4 | 3.33 |
| chr5:107216257-107218023 | 0.77 | - | Tgfbr3 | 1.37 |
| chr5:104021158-104022631 | 1.000 | - | Hsd17b11 | 3.89 |
| chr1:118647742-118649473 | 1.11 | DR5 | Tfcp2l1 | 2.77 |

**Supplemental Table S2.** Comparison of *Aldh1a2-/-* and wild-type E8.5 trunk tissue for H3K27me3 ChIP-seq and RNA-seq results to identify RA-regulated H3K27me3 ChIP-seq peaks near genes with RA-regulated expression.

| H3K27me3 ChIP-seq differential peak for *Aldh1a2* KO vs WT (mm10) | log2 fold change: H3K27me3 ChIP-seq for *Aldh1a2* KO vs WT | RARE: based on Homer TFBS analysis | nearby gene with altered expression in *Aldh1a2* KO vs WT | log2 fold change for nearby gene: RNA-seq for *Aldh1a2* KO vs WT |
| --- | --- | --- | --- | --- |
| chr18:38598986-38601292 | -1.20 | - | Spry4 | 3.43 |
| chr17:15533901-15538178 | -0.95 | - | Pdcd2 | 2.22 |
| chr17:15533901-15538178 | -0.95 | - | Tbp | 2.94 |
| chrX:104569467-104572914 | -0.90 | - | Zdhhc15 | 1.5 |
| chrX:104546302-104549188 | -0.89 | - | Zdhhc15 | 1.5 |
| chr2:118901989-118904591 | -0.85 | - | Bahd1 | 1.12 |
| chr4:129226495-129228276 | -0.85 | - | C77080 | 0.98 |
| chr4:129221927-129223300 | -0.77 | - | C77080 | 0.98 |
| chr5:15980910-15984533 | -0.75 | - | Cacna2d1 | 1.01 |
| chr3:89278454-89282169 | -0.72 | - | Efna1 | 1.23 |
| chr11:103110581-103113995 | -0.67 | - | Acbd4 | 6.49 |
| chr4:129246599-129252553 | -0.64 | DR5 | C77080 | 0.98 |
| chr10:21991375-21993681 | -0.64 | - | Sgk1 | 1.38 |
| chr6:125360514-125365732 | -0.63 | DR2 | Tnfrsf1a | 1.39 |
| chr12:54201904-54203715 | -0.63 | - | Egln3 | 3.46 |
| chr5:147297983-147318733 | -0.63 | DR2 | Cdx2 * | 1.98 |
| chrX:104536138-104539780 | -0.62 | DR2 | Zdhhc15 | 1.50 |
| chr4:98726175-98729089 | -0.61 | - | L1td1 | 1.50 |
| chr17:29080591-29082455 | -0.59 | - | Trp53cor1 | 1.91 |
| chr11:117780323-117784425 | -0.58 | DR2 | Tmc8 | 1.33 |
| chr6:122800166-122804076 | -0.54 | - | Slc2a3 | 1.35 |
| chr10:60828600-60835032 | -0.49 | - | Unc5b | 3.58 |
| chr19:45735049-45746658 | -0.49 | DR2 | Fgf8 * | 5.24 |
| chrX:94129800-94133754 | -0.48 | - | Zfx | 1.40 |
| chr2:119235078-119238967 | -0.47 | - | Spint1 | 2.74 |
| chr13:114456076-114460873 | -0.47 | DR2 | Fst * | 1.15 |
| chr19:11816637-11820013 | -0.47 | - | Stx3 | 0.90 |
| chr6:72232803-72239355 | 0.50 | - | Atoh8 | -2.59 |
| chr19:4709301-4715701 | 0.51 | - | Sptbn2 | -1.59 |
| chr3:5219339-5224436 | 0.52 | - | Zfhx4 * | -2.26 |
| chr7:130260543-130263682 | 0.56 | - | Fgfr2 | -3.06 |
| chr17:56471489-56479605 | 0.57 | DR1 | Ptprs * | -3.31 |
| chr4:144893360-144895562 | 0.59 | - | Dhrs3 | -1.11 |
| chr2:116072251-116077455 | 0.61 | DR5 | Meis2 * | -1.10 |
| chr7:70356085-70361002 | 0.63 | DR1 | Nr2f2 * | -2.32 |
| chr15:98621217-98623590 | 0.65 | - | Cacnb3 | -2.73 |
| chr1:59473538-59476300 | 0.66 | DR1 | Fzd7 | -1.64 |
| chr17:47915056-47917877 | 0.67 | - | Foxp4 * | -1.02 |
| chr10:8548515-8549917 | 0.72 | - | Ust | -1.40 |
| chr6:52156115-52158253 | 0.73 | DR5, DR2 | Hoxa1 | -5.43 |
| chr7:96211108-96212622 | 0.75 | - | Tenm4 | -2.65 |
| chrX:162887815-162889313 | 0.76 | - | Syap1 | -1.55 |
| chr11:19015536-19017169 | 0.78 | DR1 | Meis1 * | -2.64 |
| chr18:58208120-58210286 | 0.81 | - | Fbn2 | -1.65 |
| chr14:21983733-21987831 | 0.85 | - | Zfp503 | -2.68 |
| chr11:19007512-19012358 | 0.87 | DR2 | Meis1 * | -2.64 |
| chr14:16574377-16578138 | 1.02 | DR5, DR1 | Rarb * | -1.64 |

**Supplemental Table S3.**
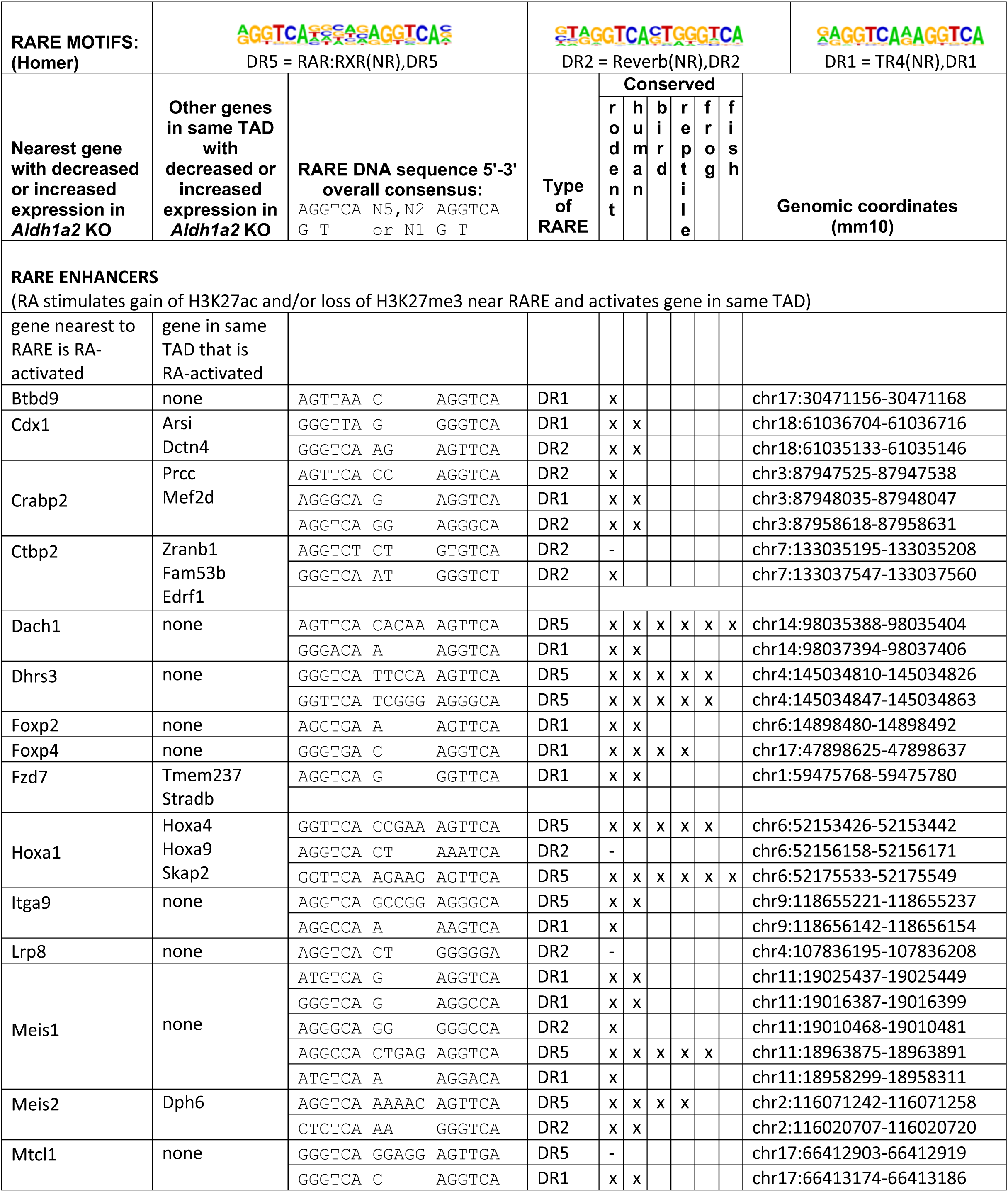
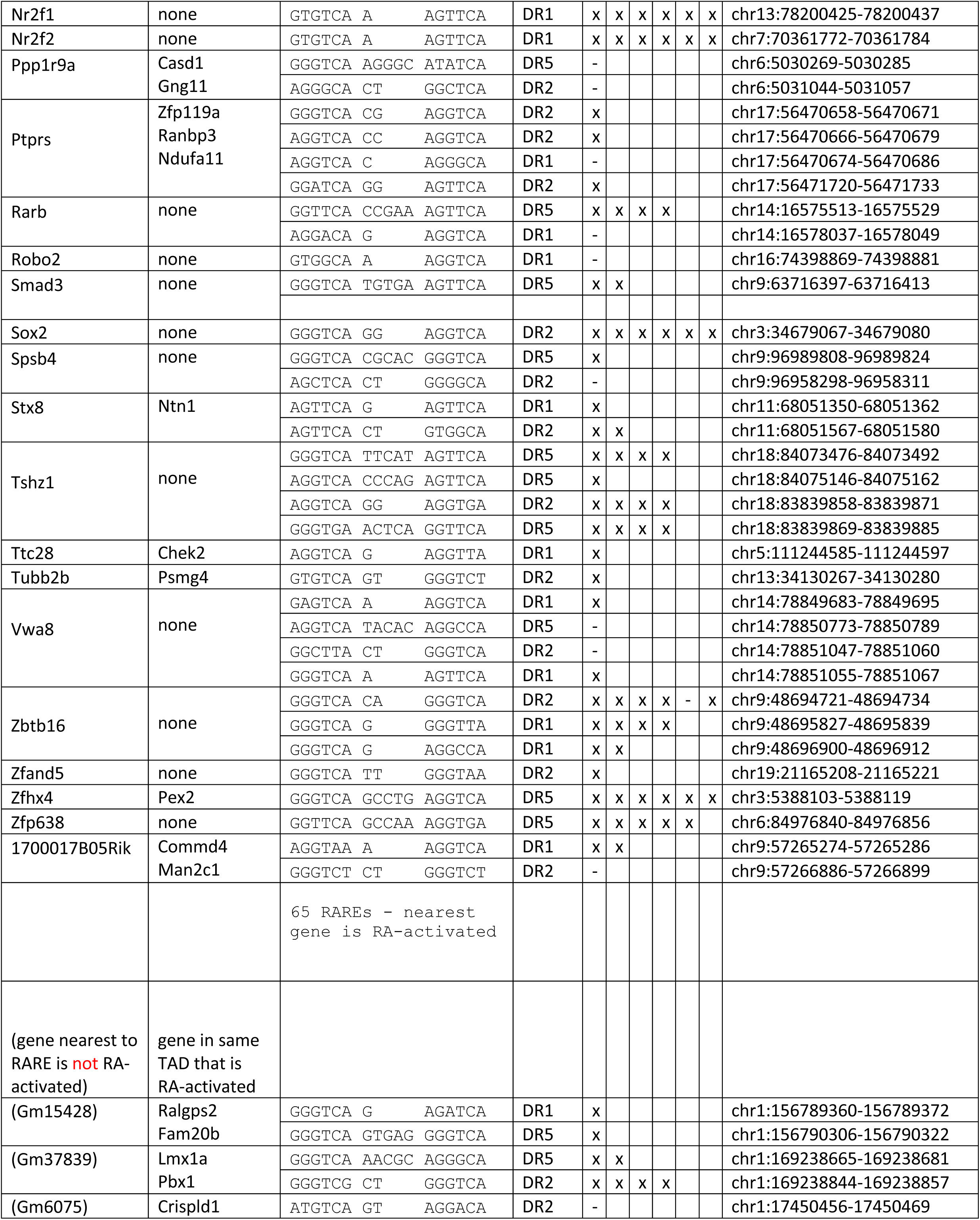
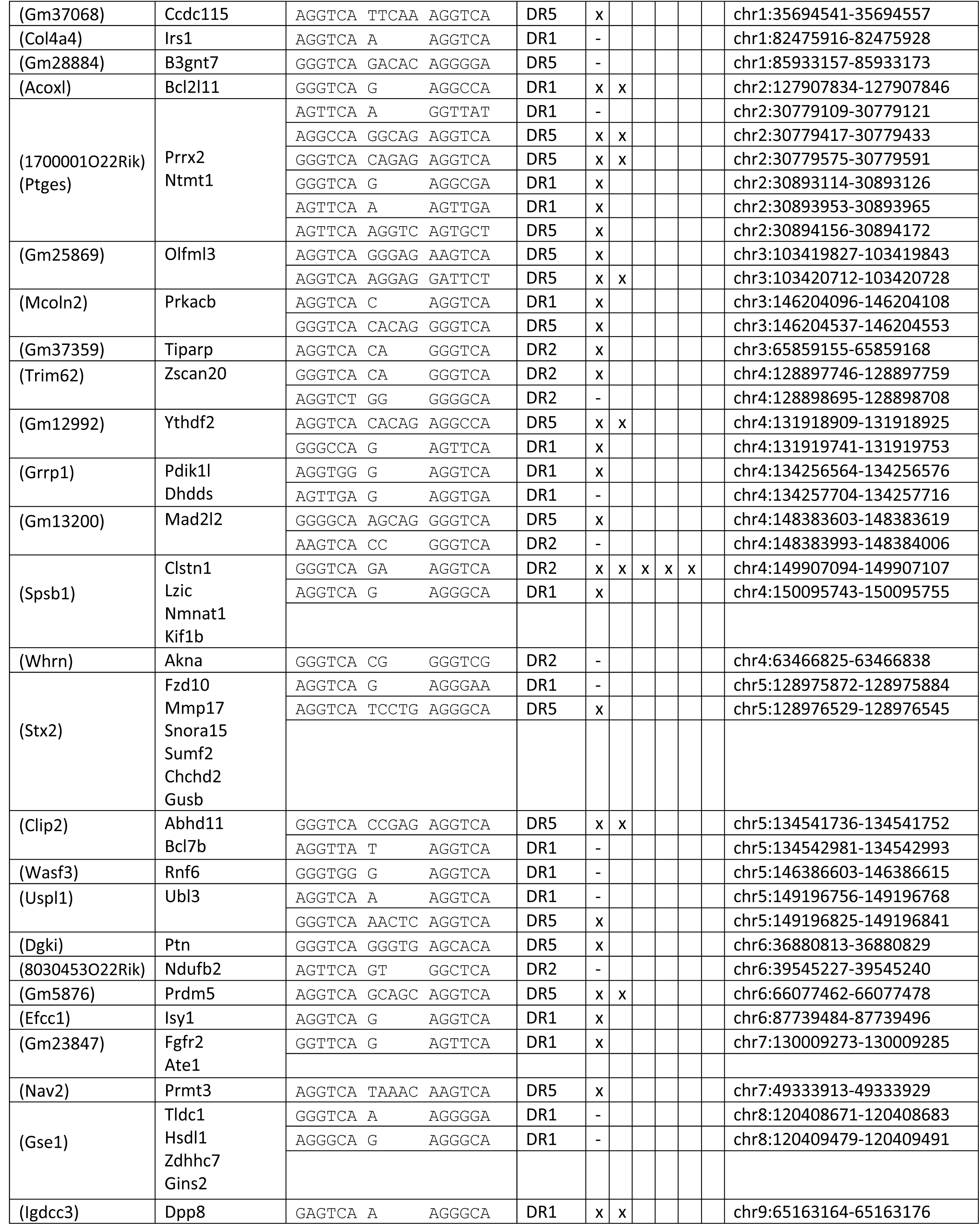
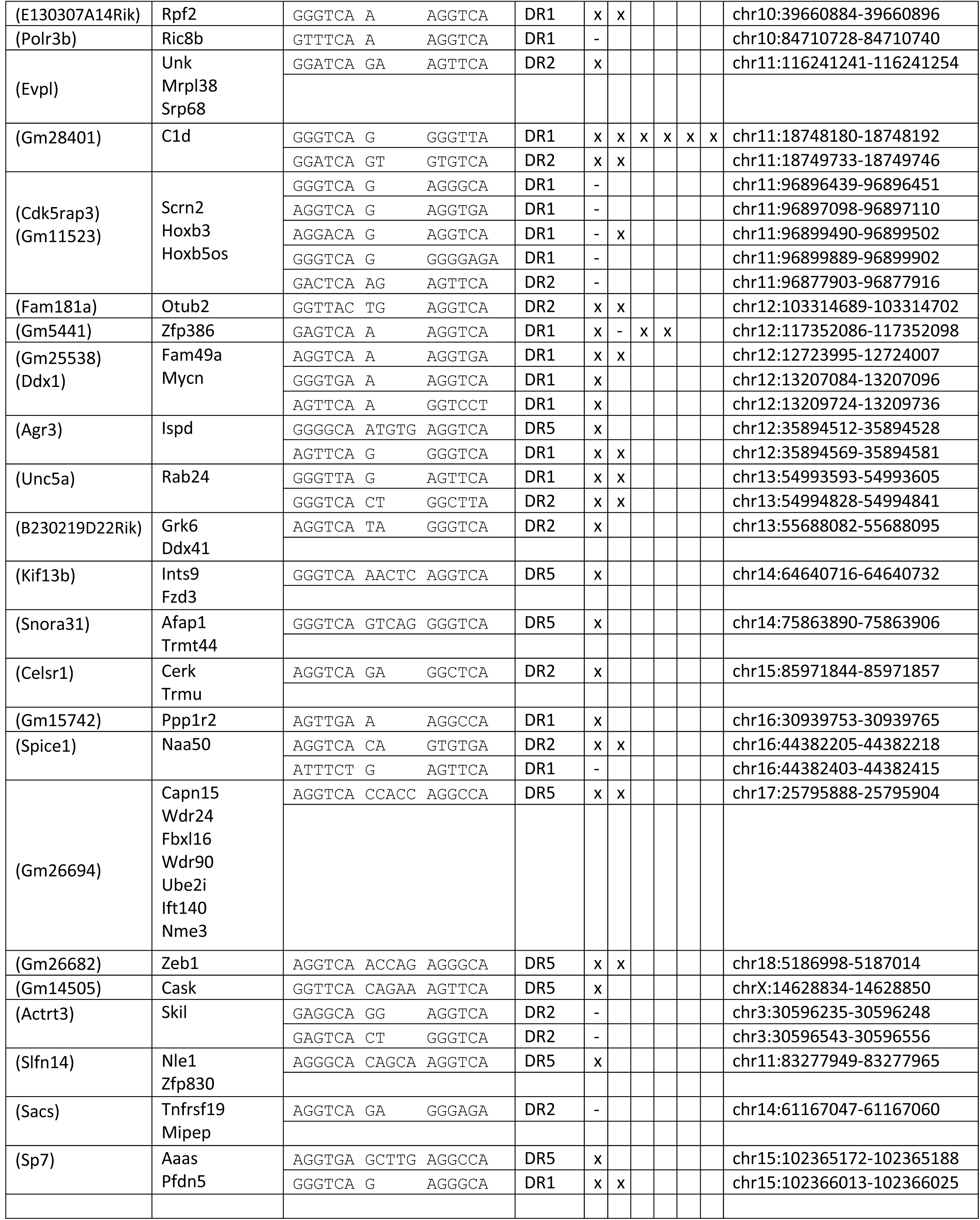
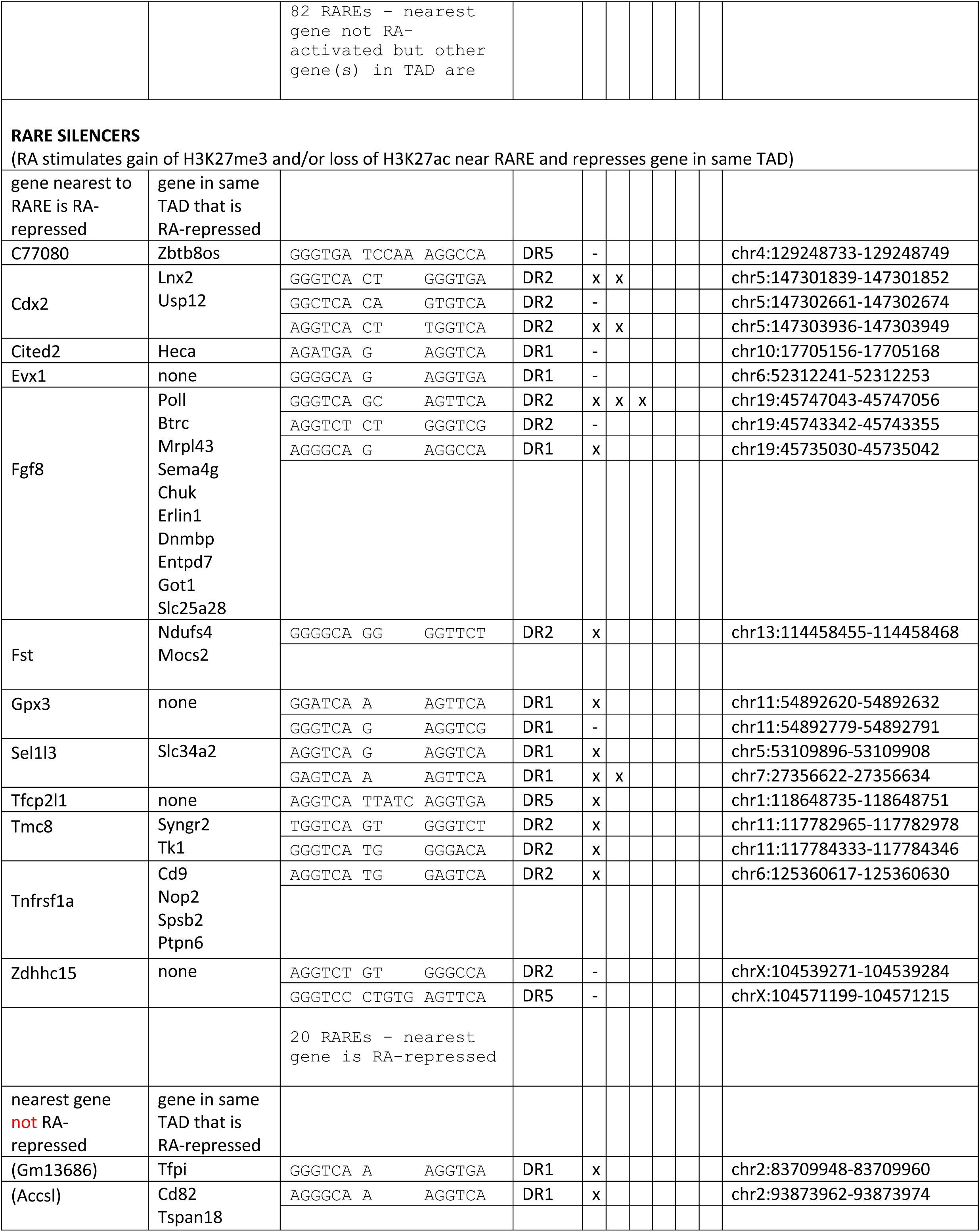
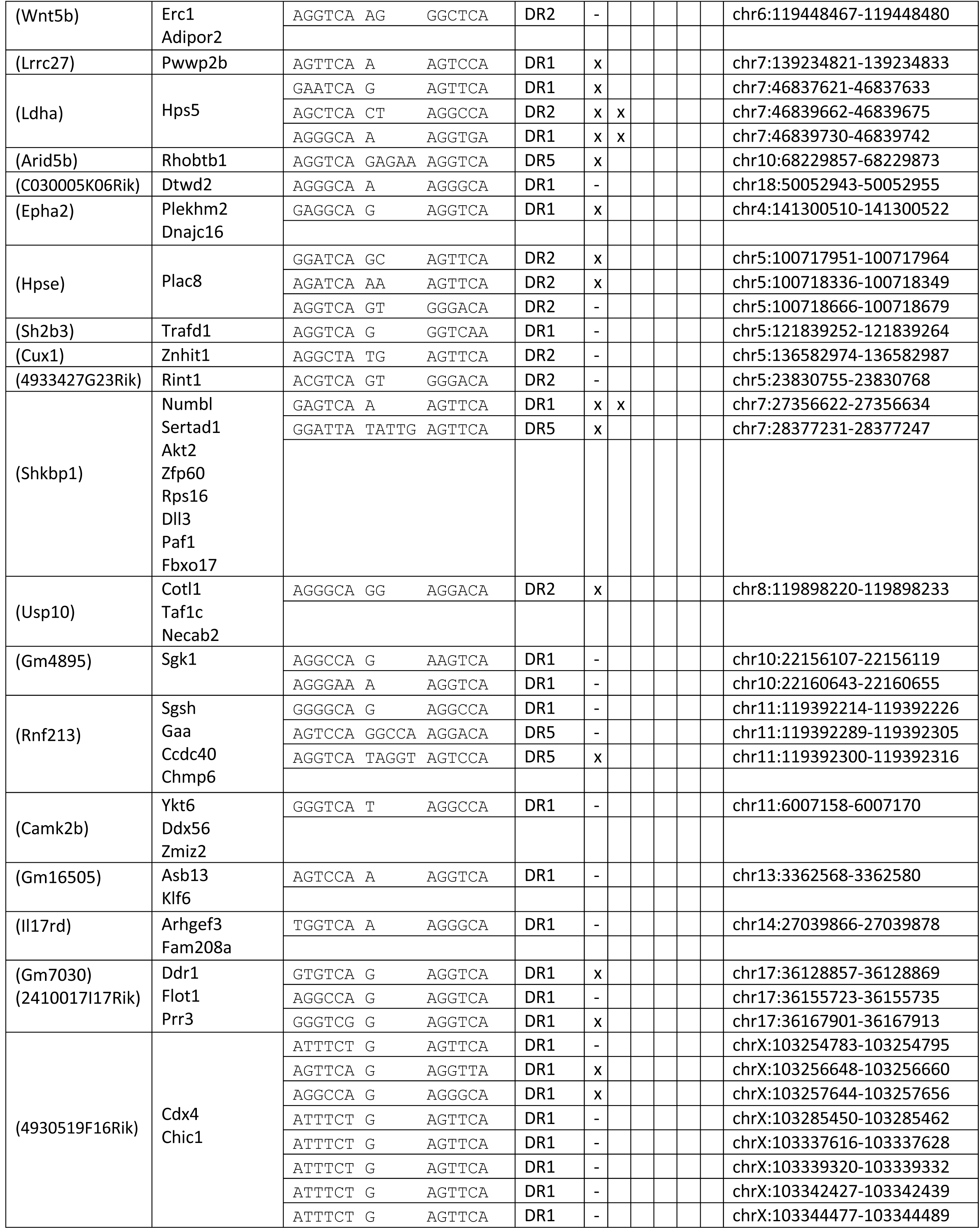
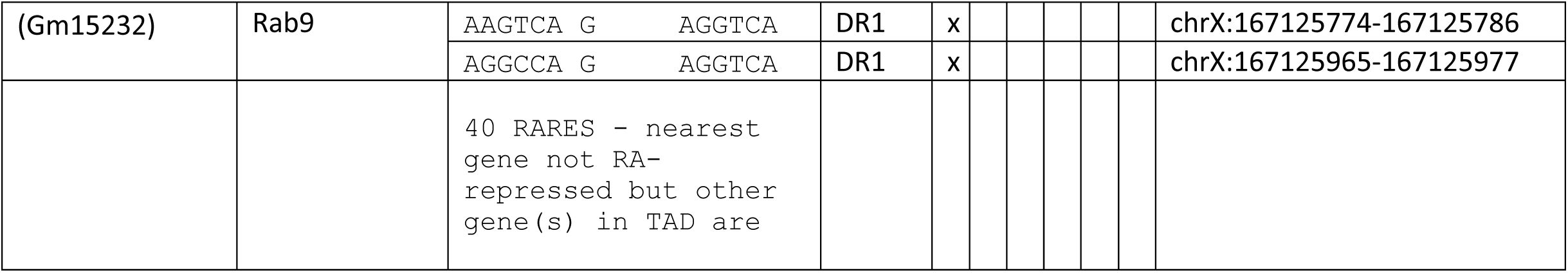
DNA sequences of all RAREs located in RA-regulated ChIP-seq peaks for H3K27ac or H3K27me3 near all RA-regulated genes in same TAD. RAREs contain no more than two mismatches to Homer consensus DR5, DR2, or DR1 RARE motifs shown here; DR, direct repeat.

**Supplemental Table S4.**
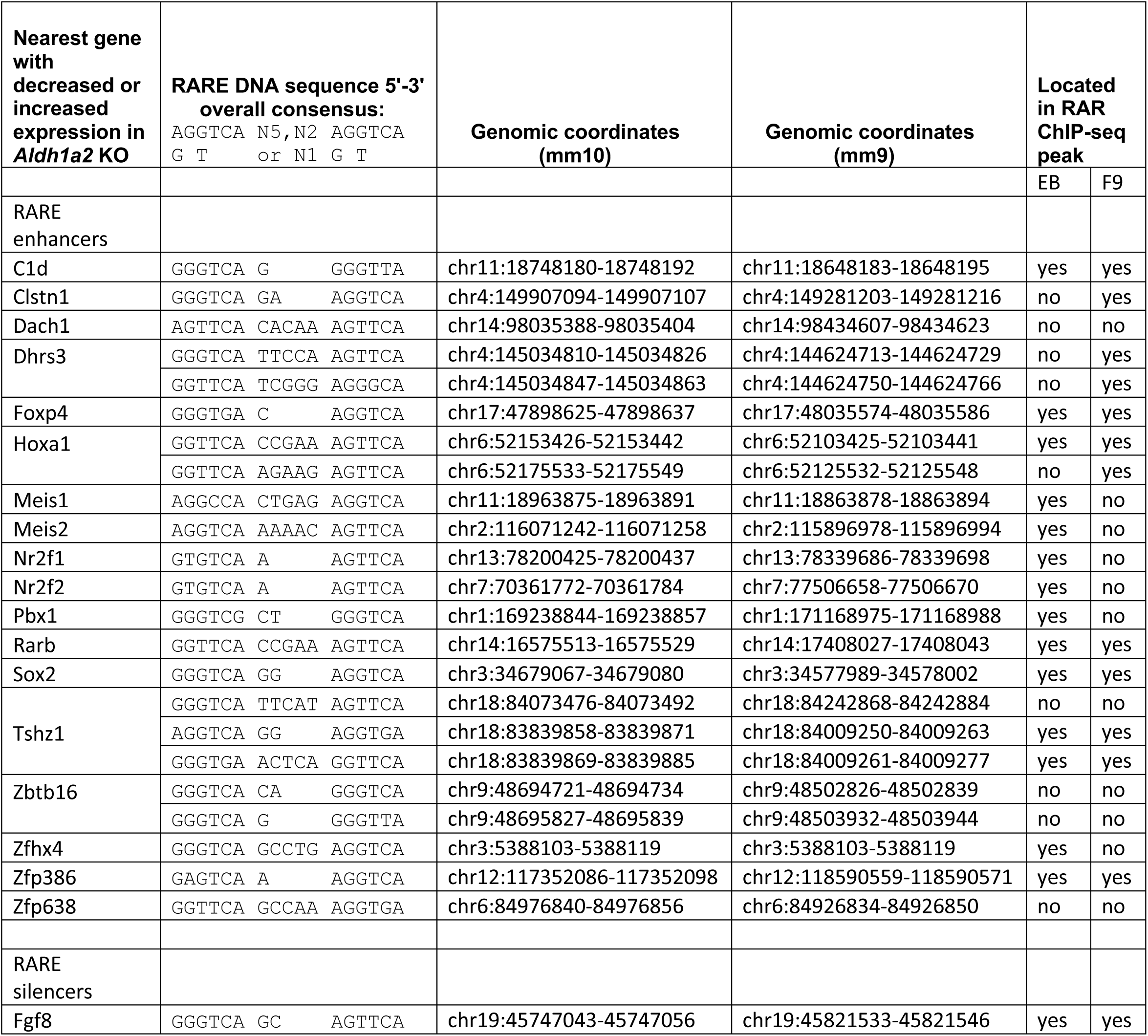
DNA sequences of highly conserved RAREs and relationship to panRAR ChIP-seq peaks for mouse embryoid bodies (EB) (Moutier et al., 2012) and F9 embryonal carcinoma cells (Chatagnon et al., 2015) reported using mm9 genomic coordinates.

